# The interaction of p130Cas with PKN3 promotes malignant growth

**DOI:** 10.1101/334425

**Authors:** Jakub Gemperle, Michal Dibus, Lenka Koudelková, Daniel Rosel, Jan Brábek

## Abstract

Protein p130Cas constitutes an adaptor protein mainly involved in integrin signaling downstream of Src kinase. Owing to its modular structure, p130Cas acts as a general regulator of cancer cell growth and invasiveness induced by different oncogenes. However, other mechanisms of p130Cas signaling leading to malignant progression are poorly understood. Here, we show a novel interaction of p130Cas with Ser/Thr kinase PKN3, which is implicated in prostate and breast cancer growth downstream of phosphoinositide 3-kinase. This direct interaction is mediated by the p130Cas SH3 domain and the centrally located PKN3 polyproline sequence. PKN3 is the first identified Ser/Thr kinase to bind and phosphorylate p130Cas and to colocalize with p130Cas in cell structures that have a pro-invasive function. Moreover, the PKN3-p130Cas interaction is important for mouse embryonic fibroblast growth and invasiveness independent of Src transformation, indicating a distinct mechanism from that previously characterized for p130Cas. Together, our results suggest that the PKN3-p130Cas complex may represent an attractive therapeutic target in late-stage malignancies.

**Summary:** Gemperle et al. present the first report of an interaction between p130Cas with the serine/threonine kinase PKN3, implicated in prostate and breast cancer growth. They show that p130Cas colocalizes with PKN3 in cell structures that have a pro-invasive function and enhance our understanding of PKN3-mediated signaling and tumor growth.

## Introduction

p130Cas (Crk associated substrate, CAS) is a molecular scaffold involved in the regulation of several processes, such as cell survival, migration, invasivity, and proliferation in both normal and pathological cells (Cabodi, del Pilar Camacho-Leal, et al., 2010; Tikhmyanova et al., 2010). Owing to its modular structure, p130Cas plays a crucial role in signaling originating from many mutated or amplified oncogenes (Nikonova et al., 2014; Tornillo et al., 2014; Tikhmyanova et al., 2010). It has been reported that knockdown of p130Cas leads to proliferative arrest in breast cancer cell lines carrying oncogenic mutations in *BRAF, KRAS, PTEN,* or *PIK3CA* (Pylayeva et al., 2009). Moreover, its involvement in Src-mediated tumorigenesis has been clearly demonstrated. For example, Src-transformed mouse embryonic fibroblasts (MEFs) exhibit increased capability to invade through Matrigel and induce metastases in mice in p130Cas-dependent manner (Honda et al., 1998; Brábek et al., 2004, 2005). Other studies in vivo have shown that p130Cas also drives the growth, aggressiveness, and progression of ErbB2-overexpressing breast tumors, including metastatic colonization of the lungs (Cabodi et al., 2010b, 2006). Correspondingly, elevated expression of p130Cas in human patients is associated with early disease recurrence and poor prognosis in several cancer types including lung, prostate, pancreas, ovarian, and mammary cancers (Fromont and Cussenot, 2011; Fromont et al., 2007; Defilippi et al., 2006; Nick et al., 2011; Tikhmyanova et al., 2010; Nikonova et al., 2014; Cabodi et al., 2010a) and is associated with hormone deprivation-mediated resistance to anti-tumor drugs (standard therapeutics) such as adriamycin (doxorubicin) and tamoxifen (Ta et al., 2008; Dorssers et al., 2001). Taken together, such evidence clearly underlines a role for p130Cas as a general regulator of cancer cell growth and metastasis as induced by different oncogenes.

The structure of p130Cas consists of an N-terminal SRC homology 3 (SH3) domain, substrate domain (SD), and serine-rich domain (SRD) followed by Src and PI3K binding regions and terminated by a CAS-family C-terminal homology domain (CCH) (Cabodi, del Pilar Camacho-Leal, et al., 2010). The majority of known p130Cas downstream signaling is attributed to tyrosine phosphorylation of a repeated YXXP motif within the p130Cas SD domain (Cabodi et al., 2010a; Defilippi et al., 2006). The level of p130Cas tyrosine phosphorylation is mainly dependent on the binding capacity of its SH3 domain, which facilitates direct interaction of p130Cas with the polyproline motif of various phosphatases (e.g., PTP1B, PTP-PEST) and kinases (FAK, PYK2) or mediates indirect association with Src via a FAK (PYK2) bridge (Fonseca et al., 2004; Ruest et al., 2001; Astier et al., 1997). Specifically, association of p130Cas via the p130Cas SH3 domain with FAK and Src at focal adhesions transmits signals that induce lamellipodia and cell migration, support cell proliferation and cell invasiveness, and block anoikis (Donato et al., 2010; Tazaki et al., 2008; Ruest et al., 2001; Nikonova et al., 2014; Defilippi et al., 2006; Brábek et al., 2005). p130Cas has also been shown to be phosphorylated at serine residues, which correlates with an invasive cell phenotype and is partially dependent on the p130Cas SH3 domain; however, responsible serine/threonine kinases have not yet been identified (Makkinje et al., 2009). In addition, p130Cas has been shown to interact with 14-3-3 proteins in a phosphoserine-dependent manner, which occurs mainly at lamellipodia during integrin-mediated cell attachment to the extracellular matrix (ECM) (Garcia-Guzman et al., 1999).

In a screen for new interaction partners of the p130Cas SH3 domain, we have recently predicted a list of candidates and verified p130Cas SH3 binding to the polyproline motifs of GLIS2 and DOK7 (Gemperle et al., 2017). Among other predicted candidates, serine/threonine PKN3 kinase, with the polyproline motif P_500_**P**P**KP**P**R**L, constitutes a member of the PKN family, which is part of the protein kinase C (PKC) superfamily of serine/threonine kinases. The role of PKN3 in tumorigenesis was identified in early reports, which showed that *PKN3* mRNA is scarce in normal human adult tissues but abundantly expressed in numerous cancer cell lines (Oishi et al., 1999). In contrast, the other members of the PKN family, PKN1 and PKN2, exhibit ubiquitous expression in human and rat tissues (Hashimoto et al., 1998; Mukai and Ono, 1994; Quilliam et al., 1996). PKN3, but not PKN1 or PKN2, has been shown to regulate malignant prostate cell growth downstream of activated phosphoinositide 3-kinase (PI3K) independent of Akt (Leenders et al., 2004) and, moreover, only PKN3 possess the polyproline sequence in its central portion that we have predicted as a potential p130Cas SH3 domain binding site (Gemperle et al., 2017). In addition, when stimulated in a fatty-acid dependent manner, the catalytic activity of PKN3 was less responsive in comparison to PKN1 and PKN2, thereby highlighting the differences in function of PKN isoforms, as well as their regulation (Oishi et al., 1999).

Using orthotopic mouse tumor models, the effect of PKN3 on cancer growth was shown by conditional reduction of PKN3 expression in tumors. In all cases, downregulation of PKN3 protein impaired primary prostate and breast tumor growth and blocked metastasis (Unsal-Kacmaz et al., 2011; Leenders et al., 2004). Correspondingly, overexpression of exogenous PKN3 in breast cancer cells further increased their malignant behavior in vitro (Unsal-Kacmaz et al., 2011). Although PKN3 KO mice appeared indistinguishable from their WT counterparts, this model also indicated the role of host stromal PKN3 in tumor progression (Mukai et al., 2016). Stromal PKN3 is enriched in primary endothelial cells as well as in osteoclasts (Uehara et al., 2017), being apart from tumor cells among the few normal cell types with significant amount of PKN3; this is consistent with the usually invasive features of endothelial cells in particular (Aleku et al., 2008). Accordingly, systemic administration of siRNA-lipoplex (Atu027) directed against PKN3 and targeting mainly the stromal compartment rather than the pathologically defined tumor entity, prevented lung metastasis in lung experimental and spontaneous breast metastasis models in a dose-dependent manner (Santel et al., 2010).

In mammalian tissues, the PKN family underlies some of the main Rho GTPase-associated protein kinase activities (Mellor et al., 1998; Vincent and Settleman, 1997; Lim et al., 2008). Specifically, Unsal-Kacmaz et al. demonstrated that PKN3 physically interacts with Rho-family GTPases, and preferentially with RhoC, a known mediator of tumor invasion and metastasis in epithelial cancers (Unsal-Kacmaz et al., 2011). However, additional molecular mechanisms by which PKN3 contributes to malignant growth and tumorigenesis are not well understood.

In this study, we have shown that PKN3 directly interacts with p130Cas and phosphorylates it in vitro and potentially in vivo. Furthermore, we have demonstrated the importance of the PKN3–p130Cas interaction for PKN3-stimulated cell growth and invasiveness in vitro and tumor growth in vivo.

## Results

### p130Cas directly interacts with PKN3

To confirm the predicted PKN3-p130Cas interaction, we first analyzed the potential of p130Cas SH3 domain variants to pull-down PKN3. The scheme of p130Cas and PKN3 mutagenesis is shown in Fig. 1 A. As predicted, only the p130Cas SH3 WT, but not phosphomimicking mutant variant (Y12E), showed strong association with PKN3 WT. Correspondingly, p130Cas SH3 WT was not able to effectively pull-down a PKN3 variant in which the target polyproline motif was mutated to P_500_APSAPRL (PKN3 mPR; Fig. 1 B; 10–50× decrease compared to WT; *p* = 0.003). Notably, a kinase-inactivating mutation in the catalytic domain of mouse PKN3 (KD; K577E; Figs. 1 A and S1 A) also caused significant decrease (4–20×; *p* = 0.007) of PKN3-p130Cas SH3 interaction (Fig. 1 B). Subsequently, we confirmed p130Cas - PKN3 interaction using co-immunoprecipitation analysis. We demonstrated that PKN3 co-precipitates with immunoprecipitated GFP-p130Cas WT and the non-phosphorylatable mutant (Y12F) (Fig. 1 C), but not with the phosphomimicking mutant (Y12E), and that p130Cas co-precipitates strongly with immunoprecipitated PKN3 WT, less with PKN3 KD, and almost not at all with PKN3 mPR (Fig. 1 D). Finally, using far-western experiments, we confirmed that the p130Cas-PKN3 interaction is direct (Fig. 1, E and F).

**Figure 1.**
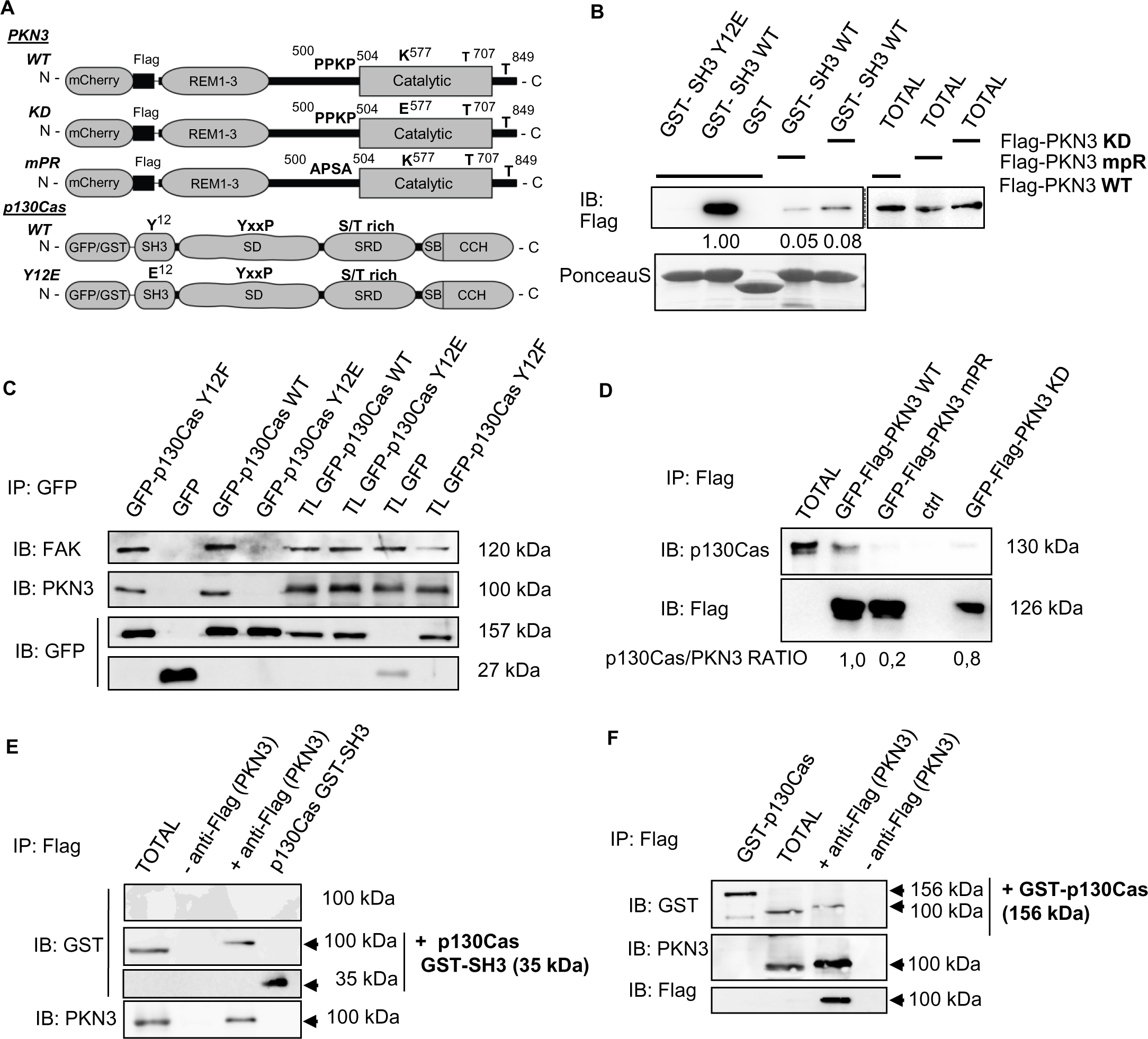
p130Cas directly interacts with PKN3 **A)** Schematic representation of PKN3 and p130Cas domains, important residues and their mutagenesis. **B)** Purified GST-fused p130Cas SH3 variants (WT, Y12E) were used to pull-down Flag-PKN3 variants (WT, mPR, KD), expressed in mouse embryonic fibroblasts (MEFs). Pulled-down proteins were immunoblotted with anti-Flag antibody, and GST-SH3 domains were stained with Ponceau-S. GST with lysate from Flag-PKN3 WT was used as a negative control. **C-D)** Binding of PKN3 to full-length p130Cas was verified by co-immunoprecipitations. **C)** The MDA-MB-231 cells were transiently transfected with indicated mouse GFP-p130Cas variants or GFP alone followed by immunoprecipitations using anti-GFP antibody. Co-immunoprecipitated PKN3 and FAK (as a positive control) were detected using anti-PKN3 antibody or anti-FAK, respectively. **D)** The MDA-MB-231 cells were transiently transfected with mouse Flag and GFP-fused PKN3 variants (WT, mPR, KD) followed by immunoprecipitations using anti-Flag sepharose and co-immunoprecipitated p130Cas was detected by anti-p130Cas antibody. In far western experiments **E-F)** Flag-PKN3 was immunoprecipitated from transfected MDA-MB-231 cells, transferred to nitrocellulose membrane and incubated with **E)** recombinant GST-p130Cas SH3 domain or **F)** whole GST-p130Cas followed by detection with anti-GST antibody. The membrane was then stripped and both endogenous (endo PKN3) as well as exogenous PKN3 (exo Flag-PKN3) were detected by anti-PKN3 and anti-Flag antibody, respectively. The upper membrane in fig. E was not incubated with recombinant GST-p130Cas SH3 domain and represents a negative control for anti GST antibody. As a positive control for GST cross-reactivity a purified GST-p13Cas SH3 in **E)** or whole GST-p130Cas in **F)** was ran alongside. TL or TOTAL: total cell lysate; IP: immunoprecipitation; Ctrl: control samples prepared from untransfected MDA-MB-231 cells.

### PKN3 colocalizes with p130Cas in lamellipodia and podosome rosettes

To further assess the p130Cas-PKN3 interaction, we analyzed the co-localization of p130Cas and PKN3 in cells. Specifically, we analyzed the dynamic localization of p130Cas and PKN3 in p130Cas−/− MEFs co-expressing GFP-p130Cas, variants of mCherry-PKN3, and CFP-LifeAct. We found that both p130Cas and PKN3 were enriched in lamellipodia of MEFs (Fig. 2 A; Movies S1 and 2). Localization of the PKN3-mPR mutant to lamellipodia was slightly but significantly impaired (Fig. 2, A and B), suggesting that the polyproline sequence of PKN3 is important for its targeting to lamellipodia. However, cells expressing GFP-Flag-PKN3 KD exhibited more pronounced filopodia-like protrusions and the presence of lamellipodia was extremely rare compared to that in cells expressing other variants of PKN3, preventing quantification of PKN3 membrane localization (Fig. S1 B and Movie S3).

**Figure 2.**
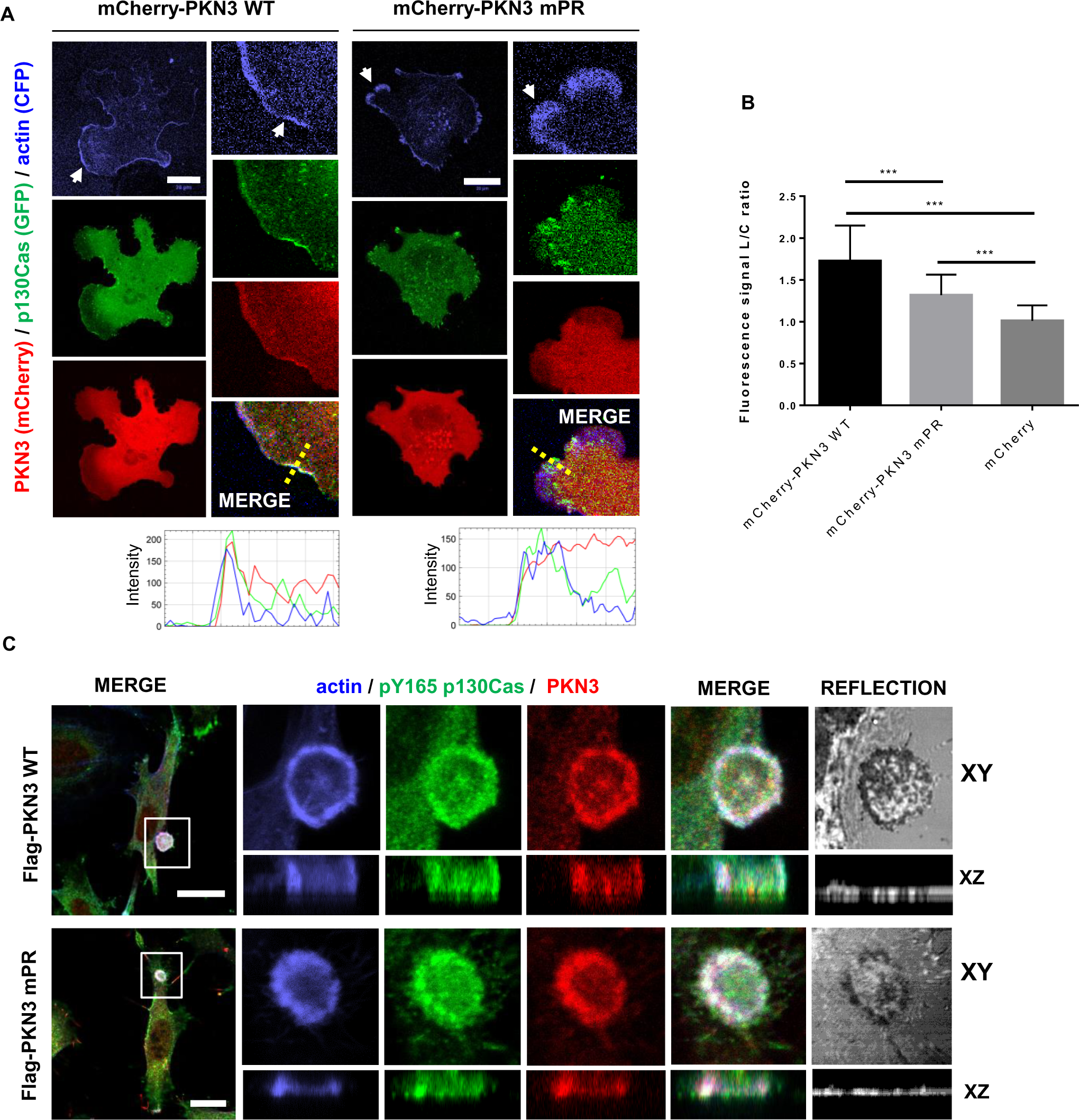
PKN3 colocalizes with p130Cas in lamellipodia and podosome rosettes. Representative images are shown. **A)** p130Cas−/− MEFs plated on fibronectin (FN) were transfected by GFP-p130Cas, CFP-LifeAct and mCherry-PKN3WT or mCherry-PKN3 mPR and imaged live 24 h after transfection. White arrow indicates lamellipodia. Histogram of dotted straight line is shown. **B)** Quantification of mCherry-PKN3 WT, mCherry-PKN3 mPR and mCherry localization to lamellipodia (LifeAct as marker) was calculated as described in methods (values are mean ± SD from three independent experiments, n>50 measurements – 3 per cell; *P<0,001, one-way ANOVA on ranks followed by Dunn’s post-hoc test). **C)** Src-transformed p130Cas−/− MEFs co-expressing p130Cas (SC) and mouse Flag tagged PKN3 WT or Flag-PKN3 mPR are shown. Cells were grown on FN-coated coverslips for 48 h, fixed and stained for p130Cas by anti-pTyr165 p130Cas antibody (pY165 p130Cas; 2nd 405), for actin by Phalloidin 488 and for Flag-PKN3 by anti-Flag antibody (2nd 633). Reflection (670 nm) indicates fibronectin degradation. All scale bars represent 20 µm. Cell were imaged by Leica TCS SP8 microscope system equipped with Leica 63×/1.45 oil objective.

PKN3 has been recently shown to localize to specific actin-rich structures termed podosome rings and belts in osteoclasts (Uehara et al., 2017). Formation of similar structures, termed podosome rosettes, can be induced in MEFs when transformed by activated Src (SrcF) (Tarone et al., 1985). Notably, p130Cas was shown to be critical for their formation (Brábek et al., 2004). To investigate whether PKN3 localizes to podosome rosettes, we expressed the Flag-PKN3 variants in SrcF-transformed p130Cas−/− MEFs re-expressing p130Cas (SC cells) and analyzed their localization using confocal microscopy on fixed cells. Both PKN3 WT and mPR were enriched in podosome rosettes and co-localized there along with p130Cas and actin, suggesting that p130Cas is not responsible for PKN3 targeting to podosome rosettes (Fig. 2 C).

### PKN3 activity is important for stress fiber formation

In live cell imaging/microscopy experiments, we noticed temporary co-localization of GFP-PKN3 WT or mPR with stress fibers (mCherry-LifeAct) (Fig. S1, B and C). As PKN3 downregulation in HUVEC cells leads to disruption of stress fiber formation (Mopert et al., 2012) and p130Cas expression has been previously shown to affect stress fiber morphology (Honda et al., 1998), we analyzed the importance of PKN3 activity and interaction with p130Cas on stress fiber formation. Therefore, we co-transfected p130Cas-GFP WT and mCherry-Flag-fused PKN3 variants (WT, mPR, KD) or mCherry to p130Cas−/− MEFs (Fig. 3 A). Cells expressing PKN3 WT or mPR displayed prominent stress fibers, suggesting that interaction between p130Cas and PKN3 is not important for stress fiber formation. In contrast, transfection of cells by PKN3 KD greatly reduced stress fiber formation, instead inducing cortical localization of F-actin (Fig. 3 A).

**Figure 3.**
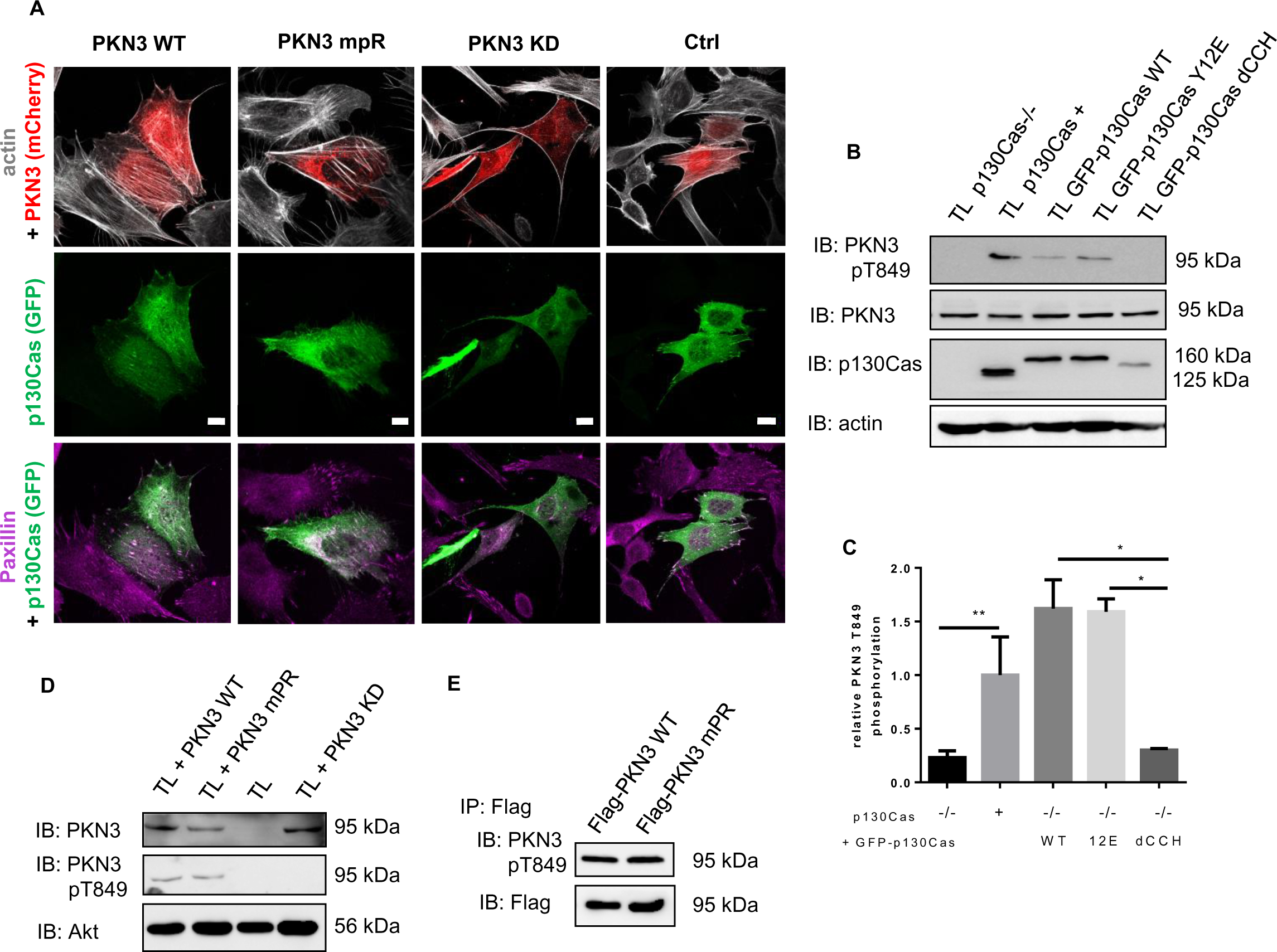
PKN3 activity is important for stress fibers formation and is stimulated by expression of p130Cas. **A)** p130Cas−/− MEFs growing on FN-coated cover slips were co-transfected by GFP-p130Cas and mCherry-PKN3 fusion variant (WT, mPR, KD) or mCherry. After 48 h cells were fixed and imaged by Leica TCS SP2 microscope (63×/1.45 oil objective). Stress fibers were visualized by Phalloidin (405) and focal adhesions by anti-Paxillin staining (2nd 633). Representative images are shown. Scale bars represent 20 µm. **B)** p130Cas−/− MEFs or p130Cas−/− MEFs re-expressing p130Cas or transfected by GFP-fused p130Cas variants (WT, YE, dCCH) were lysed in RIPA buffer, blotted to nitrocellulose membrane and analyzed for endogenous PKN3 activity by antibody anti-phosphoThr849 of PKN3 (pT849 PKN3). Expression of p130Cas mutants was verified by anti-p130Cas antibody and loading by anti-PKN3 and anti-actin antibody. **C)** Densitometric quantification of PKN3 activity (pT849 PKN3 phosphorylation). Error bars indicate means ± SD from three independent experiments. Statistical significance was evaluated by one-way repeated ANOVA followed by Turkey’s post-hoc test. **D)** Lysates or **E)** immunoprecipitates (by Flag-sepharose) from p130Cas−/− MEFs re-expressing p130Cas and overexpressing PKN3 variants (WT, mPR, KD) were immunoblotted by anti-PKN3, anti-pT849 PKN3 and anti-Akt antibodies (loading control).

### p130Cas stimulates PKN3 kinase phosphorylation on Thr849 independently of p130Cas-PKN3 interaction

It has previously been shown that PKN3 turn motif site phosphorylation (human Thr860, homologous to mouse Thr849) correlates with PKN3 activity and that active PKN3 is predominantly localized in the nucleus (Unsal-Kacmaz et al., 2011; Leenders et al., 2004). To determine whether p130Cas influences PKN3 activation we analyzed the activity of endogenous mouse PKN3 using an anti-pThr849 antibody in the presence or absence of p130Cas protein. PKN3 phosphorylation on Thr849 was lower in p130Cas−/− MEFs than in p130Cas−/− MEFs re-expressing p130Cas or transfected by GFP-p130Cas (Fig. 3, B and C). Expression of the Y12E variant of GFP-p130Cas, which does not bind PKN3, also increased Thr849 phosphorylation of PKN3. Consistent with these findings, the PKN3 mPR mutant, which is unable to bind p130Cas, did not differ with regard to pThr849 phosphorylation status from PKN3 WT (Fig. 3, D and E). Notably, however, Thr849 phosphorylation was not increased by expression of GFP-p130Cas without the CCH domain (dCCH) (Figs. 3 A and B, and S2 D). PKN3 KD, used as a negative control for anti-pThr849 antibody specificity, was not phosphorylated, as expected (Fig. 3 D). Taken together, the data suggest that p130Cas expression induces PKN3 activation and that this activation is independent of p130Cas–PKN3 interaction.

### p130Cas–PKN3 co-expression and p130Cas Ser428(432) and PKN3 Thr860 phosphorylation are positively correlated in human breast (prostate) tumors

Previous studies showed positive correlation between PKN3 or p130Cas/BCAR1 protein levels and cancer progression in patients with breast or prostate cancer (Strumberg et al., 2012; Schultheis et al., 2014; Dorssers et al., 2004; Leenders et al., 2004; Oishi et al., 1999; Nikonova et al., 2014; Fromont and Cussenot, 2011). To test the assumed link between PKN3 and p130Cas signaling, we further performed cross-correlation analysis of publicly available transcriptomic data using the cBio Cancer Genomics Portal (cbioportal.org) (Gao et al., 2013). We focused on invasive breast carcinoma (1100 tumors in TCGA, provisional) and prostate adenocarcinoma studies (499 tumors, TCGA, provisional). Within these sets of RNA-Seq data, we evaluated the mRNA expression of PKN3 and p130Cas/BCAR1 and ran correlation statistical analysis (Fig. 4, A-C). This analysis showed that elevated expression of *PKN3* significantly positively correlates with the elevated expression of p130Cas/BCAR1 and vice versa in both patients with breast cancer and those with prostate cancer (Figs. 4 A and S2 A). Similarly, the level of p130Cas/BCAR1 expression increased with the level of Src kinase, signaling through which is highly associated with p130Cas/BCAR1 (Brábek et al., 2004; Fonseca et al., 2004; Figs. 4 B and S2 B). In contrast, expression of *PKN2,* which does not possess a p130Cas interaction site and therefore does not suggest crosstalk with the p130Cas/BCAR1 signaling circuit, had negative correlation to mRNA levels of p130Cas/BCAR1 (Figs. 4 C and S2 C and D).

**Figure 4.**
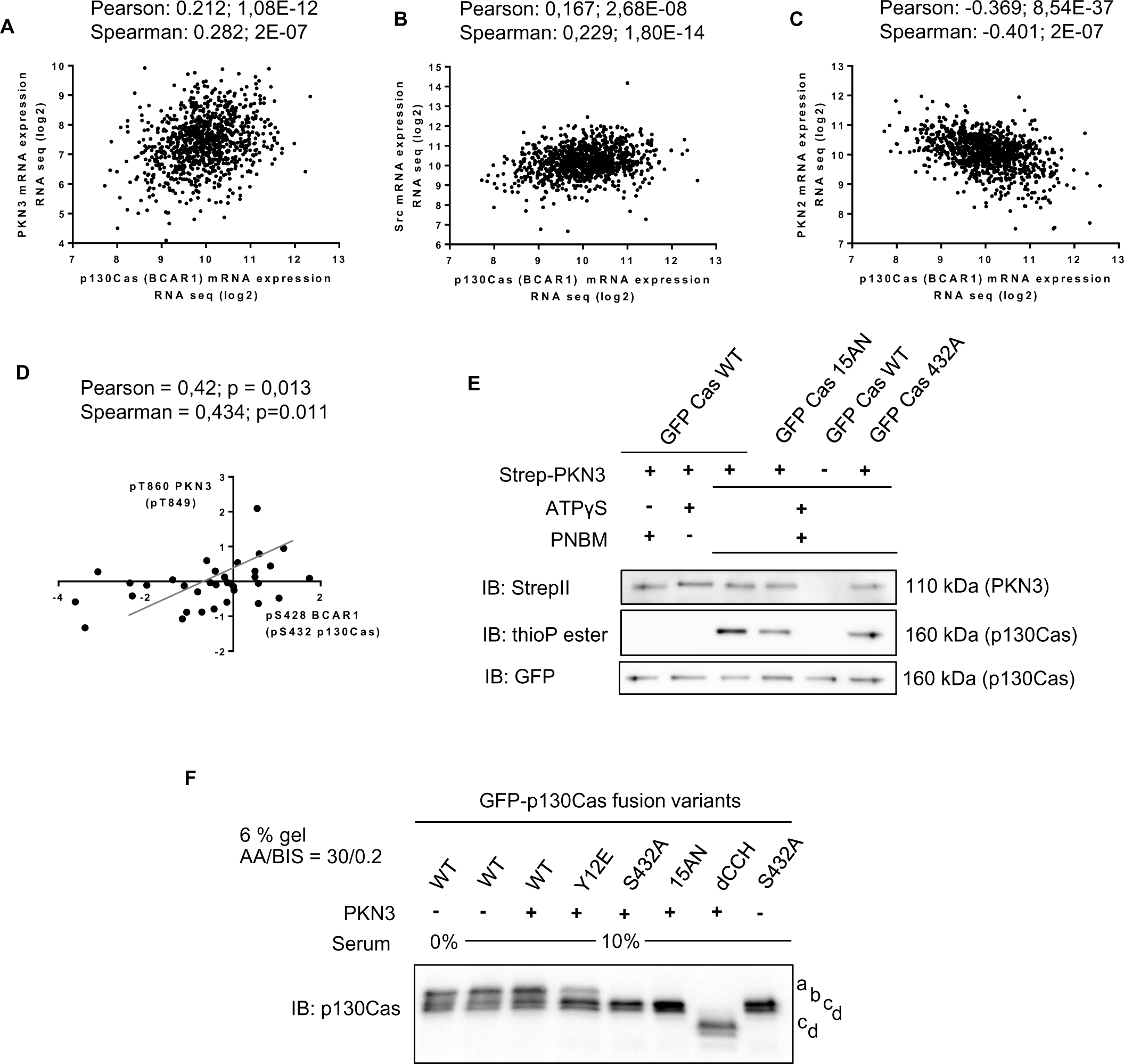
PKN3 phosphorylates p130Cas and PKN3 activity correlates with p130Cas phosphorylation in human breast carcinomas. **A-C)** Correlation statistics (graphs) of publicly available RNA seq data of co-expression of **A)** human p130Cas/BCAR1 and PKN3; **B)** p130Cas/BCAR1 and Src; **C)** p130Cas/BCAR1 and PKN2 or graph **D)** showing positive linear dependency of protein phosphorylation levels (log ratio values) of PKN3 Thr860 and p130Cas/BCAR1 Ser428 in invasive human breast tumors. **E)** Kinase reactions in vitro with precipitated Strep-PKN3 and GFP fusion p130Cas variants (shown as Cas). Reactions were carried out in the presence of ATPγS followed by alkylation with PNBM and detection with specific anti-thiophosphate esters (thioP ester) antibody. Combinations of PNBM (alkylation reagent) or ATPγS were facilitated to exclude false positive signals. Antibodies anti-StrepII and GFP were used to detect PKN3 kinase or GFP-fused p130Cas variants, respectively. Representative blots are shown. **F)** Lysates from p130Cas−/− MEFs re-expressing GFP-fused p130Cas variants with or without mCherry-PKN3 overexpression were run on SDS-PAGE using an acrylamide/bisacrylamide ratio of 30:0.2 followed by immunoblotting and detection by anti-p130Cas antibody. a-d refers to different GFP-p130Cas isoforms.

In parallel we analyzed the available data on protein phosphorylation levels of PKN3 and p130Cas/BCAR1 (Clinical Proteomic Tumor Analysis Consortium (CPTAC), MS analysis of 34 invasive breast carcinoma tumors). Statistical analysis revealed that phosphorylation of PKN3 at Thr860 (homologous to mouse Thr849), which reflects PKN3 activity, and p130Cas/BCAR1 at Ser428 (conserved homolog to mouse Ser432 and shown in cBioportal analysis as p130Cas/BCAR1 Ser474) exhibits strong positive correlation (Pearson test 0.42, *p* value 0.0134; Spearman test 0.434, *p* value 0.0106; Fig. 4 D). Taken together, the cross-correlation analysis of the transcriptomic and phosphoproteomic data showed a positive correlation between PKN3 and p130Cas/BCAR1 expression in invasive breast carcinoma and prostate adenocarcinoma tumors, and positive correlation between PKN3 activity and the level of p130Cas/BCAR1 phosphorylation on Ser428 in invasive breast carcinoma tumors.

### PKN3 phosphorylates mouse p130Cas on Ser432

p130Cas has also been previously reported to undergo serine phosphorylation under various conditions; e.g., during mitosis or cell adhesion. However the kinase responsible for the phosphorylation has not yet been identified (Garcia-Guzman et al., 1999; Makkinje et al., 2009; Yamakita et al., 1999). Our cross-correlation analysis of the transcriptomic and phosphoproteomic data showed that phosphorylation of human p130Cas/BCAR1 on Ser428 correlates with increased PKN3 activity, indicating that PKN3 might be responsible for this phosphorylation. Human Ser428 of p130Cas/BCAR1 corresponds to mouse p130Cas Ser432, which has conserved surrounding sequence (KRL**S**A) and fits well to the known PKN3 phosphorylation motif (Collazos et al., 2011). As for the majority of p130Cas potential Ser/Thr phosphorylation sites, Ser432 is localized in the p130Cas SRD domain. To test the assumed ability of PKN3 to phosphorylate p130Cas, we prepared a GFP-fused p130Cas mutant for Ser432 (S432A) and p130Cas mutated in SRD in such a manner that all 15 Ser/Thr sites were substituted with Ala or Asn (15AN). The substitution of several Ser/Thr to Asn, instead of more common substitution to Ala, was used with the aim to preserve the helical structure of the SRD. To determine whether PKN3 could phosphorylate p130Cas in the SRD domain we performed kinase assays in vitro with precipitated Strep-Flag-PKN3 and newly prepared GFP-p130Cas variants (WT, 15AN, and S432A) in the presence of ATPγS followed by detection with a specific antibody against anti-thiophosphate ester (Fig. 4 E). GFP-p130Cas WT exhibited the highest thiophosphorylation compared to GFP-p130Cas 15AN and Ser432, which were thiophosphorylated to a similar extent. Partial thiophosphorylation of GFP-p130Cas 15AN indicated that PKN3 could phosphorylate p130Cas also outside SRD domain (Fig. 4 A). Notably, PKN3 autophosphorylation was also observed (Fig. S2 E). These results verified the presence of a PKN3 phosphorylation motif in the sequence surrounding Ser432 and indicated that PKN3 phosphorylates p130Cas on Ser432 in vitro.

Protein p130Cas, similarly to Nedd9 (another member of the p130Cas family; HEF1), exists in cells in two main isoforms, which are attributed to different Ser/Thr phosphorylation (Hivert et al., 2009; Makkinje et al., 2009). Depending on the cell type, p130Cas protein is present upon SDS-PAGE as a single protein or detected as a doublet with the increased proportional representation of the upper band correlating with cell aggressive properties (Janoštiak et al., 2011; Makkinje et al., 2009). To test the biological relevance of p130Cas Ser432 phosphorylation in cells, we prepared lysates from p130Cas−/− MEFs with reintroduced GFP-p130Cas mutants with or without overexpressing mCherry-PKN3 WT and examined the p130Cas migration profile after separation on SDS-PAGE using an acrylamide/bisacrylamide ratio of 30:0.2 (Fig. 4 F), which improves the separation of Ser/Thr phosphorylation-induced gel shifts, similarly to Hivert et al. (Hivert et al., 2009). GFP-p130Cas WT migrated as 4 bands with the most pronounced form migrating the slowest. GFP-p130Cas Y12E displayed a shift of the p130Cas protein content from the slowest migrating band toward faster migrating forms. In comparison, the 15 Ser/Thr mutations (15AN) migrated only in the form of two faster bands, similarly to the single mutation of Ser432 (S432A mutant), and was not changed by PKN3 overexpression (Fig. 4 F). Furthermore, GFP-p130Cas without the CCH domain also migrated in the form of two faster bands. We concluded that p130Cas SH3 and CCH domains are important for p130Cas Ser/Thr phosphorylation and that the presence of the slowest migration form of p130Cas is associated with Ser432 phosphorylation. However, we failed to promote a switch of the slower migrating p130Cas (WT) isoform to faster ones by serum starvation (Fig. 4 F) or by maintaining the cells in suspension (Fig. S2 F); therefore, we could not demonstrate the change of p130Cas WT SDS-PAGE migration pattern by PKN3 overexpression.

### PKN3 overexpression regulates the growth of MEFs in a PKN3–p130Cas interaction-dependent manner

Both PKN3 and p130Cas have been implicated in the regulation of malignant cell growth (Cabodi et al., 2010b, 2006; Leenders et al., 2004). Although PKN3 is scarce in normal human adult tissues except in endothelial cells (HUVECs) (Aleku et al., 2008), a recent study showed that PKN3 is also present in moderate levels in MEFs, supporting these cells as a physiologically relevant model (Mukai et al., 2016). To investigate whether PKN3 regulates cell proliferation in MEFs and whether this is dependent on p130Cas, we prepared stable MEF cells with doxycycline (Dox) inducible mCherry-fused PKN3 WT or mCherry alone in the p130Cas null background with or without re-expression of p130Cas (Fig. 5 A). mCherry-PKN3 WT expression started from 5 h and reached saturation within 24 h post-treatment with Dox (Fig. 5 B). Cell morphology was unaffected by mCherry-PKN3 WT expression; however, starting 5 h after adding Dox, MEFs re-expressing p130Cas exhibited increased growth rate compared to that of non-induced controls (Fig. 5 C). Induction of mCherry alone as an additional control had no effect and the growth rate of p130Cas−/− MEFs was not affected by PKN3 expression (Fig. 5, D and E), indicating that the PKN3-mediated regulation of cell growth was p130Cas-dependent.

**Figure 5.**
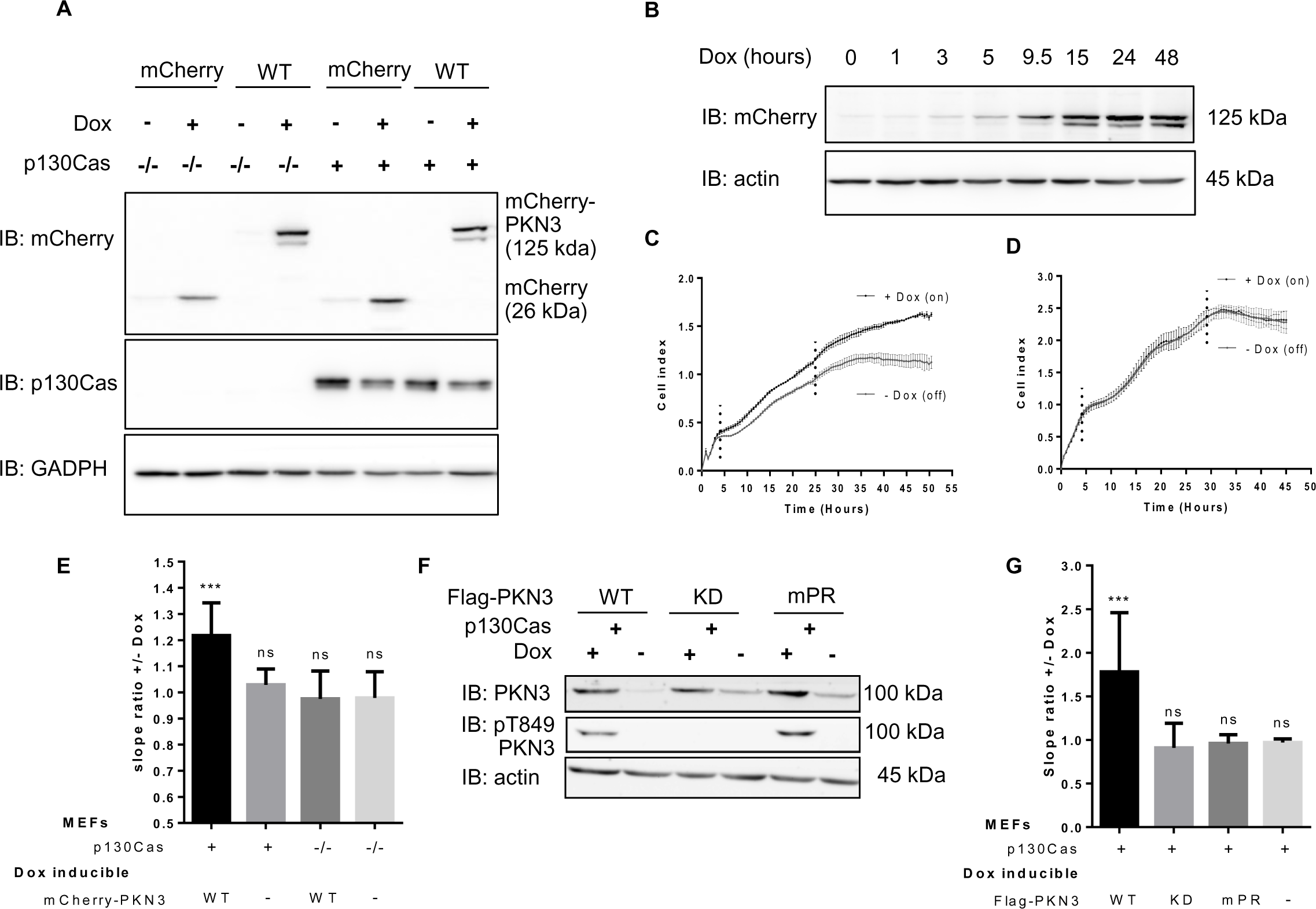
PKN3 overexpression regulates growth of MEFs and this effect requires PKN3 – p130Cas interaction. **A)** Immunoblotted lysates from MEFs p130Cas−/− or MEFs p130Cas−/− re-expressing p130Cas (p130Cas+) treated by Doxycycline (Dox) to induce expression of mCherry-PKN3 or mCherry alone. p130Cas presence was detected by ani-p130Cas antibody and mCherry epitope by anti-mCherry antibody. **B)** Dynamics of mCherry-PKN3 expression after supplementation with Dox shown by immunoblot with anti-mCherry antibody. **C-E)** Effect of induced mCherry-PKN3 expression on cell growth. Representative graphs showing growth of MEFs p130Cas/ re-expressing p130Cas (p130Cas+) **C)** or MEFs p130Cas/ **D)** measured in real-time using the xCELLigance RTCA (real-time cell analysis) system instrument. **E)** Quantification of cell growth change induced by mCherry-PKN3 expression (“-“ indicates inducible mCherry expression used as negative control). Slope ratios reflecting cell growth were calculated from the log growth phase of cell growth (indicated by dotted lines; see C and D). **F)** Immunoblotted lysates from MEFs p130Cas−/− re-expressing p130Cas (p130Cas+) treated or not treated by Dox which induced expression of Flag-fused PKN3 variants (WT, mPR, KD, empty vector). Stimulated overexpression of PKN3 was detected by anti-PKN3 antibody and its activity by antibody anti-pT849 PKN3. **G)** Quantification of cell growth change stimulated by Dox-inducible expression of Flag-fused PKN3 variants (WT, mPR, KD) in MEFs p130Cas−/− re-expressing p130Cas (p130Cas+). All error bars indicate means ± SD calculated from 3-5 independent experiments (each in triplicates). Statistical significance was always calculated between induced and non-induced cells and evaluated by one-way repeated ANOVA followed by Turkey post-hoc test (***P < 0.001).

To investigate whether PKN3-mediated induction of cell growth rate required PKN3 interaction with p130Cas, we prepared stable cell lines of p130Cas−/− MEFs re-expressing p130Cas with Dox-inducible Flag-fused PKN3 variants (WT, mPR, KD) (Fig. 5 F). Only the induction of expression of PKN3 WT, but not mPR or KD, led to the increase of cell growth compared to that of non-induced controls (Fig. 5 G). Increase of cell proliferation induced by PKN3 variants was also verified by AlamarBlue assay (Fig. S3 A) with similar outcome.

To test whether PKN3 promoted cell growth of MEFs by phosphorylation of the p130Cas SRD domain, we reintroduced GFP-p130Cas variants (WT, 15AN) or GFP as control to p130Cas−/− MEFs with Dox-inducible mCherry-PKN3 (Fig. S3 B). Both GFP-p130Cas WT and 15AN, but not GFP alone, changed cell morphology, as characterized by an increased number of cell protrusions (Fig. S3 C). This morphology was not changed by Dox-inducible mCherry-PKN3, although Dox treatment increased the cell growth of cells expressing GFP-p130Cas WT and 15AN compared to that of non-induced controls (Fig. S3 D). Therefore, the effect of PKN3 on cell growth was probably independent of p130Cas SRD domain phosphorylation.

To investigate which signaling pathways are involved in the accelerated cell growth induced by PKN3, we analyzed the activation status of STAT3, ERK, Akt, MLC, mTOR, and Src signaling (Fig. S4 A). In agreement with previous studies, we did not find any significant changes (Leenders et al., 2004). Only Src activity was slightly increased by PKN3 expression in a p130Cas-dependent manner, which is consistent with a recent study demonstrating that PKN3 could promote Src activity in osteoclasts (Fig. S4, A and B) (Uehara et al., 2017). Taken together, these results indicated that PKN3 regulates the growth of MEFs and that this effect requires PKN3-p130Cas interaction independently of p130Cas SRD domain phosphorylation.

### Interaction of p130Cas with PKN3 is required for PKN3-dependent increase of invasiveness

The localization of PKN3 in lamellipodia suggests its importance in cell migration. In p130Cas−/−MEFs re-expressing p130Cas, we analyzed the effect of inducible expression of PKN3 on 2D migration using a wound healing assay. We found that PKN3 expression has no effect on the 2D migration on either plastic or fibronectin (FN) (Fig. 6, A and B). These results are consistent with a study of Lachmann et al. (Lachmann et al., 2011) where only siRNA anti-PKN1 or anti-PKN2, but not anti-PKN3, had an effect on wound healing closure of 5637 bladder tumor cells.

**Figure 6.**
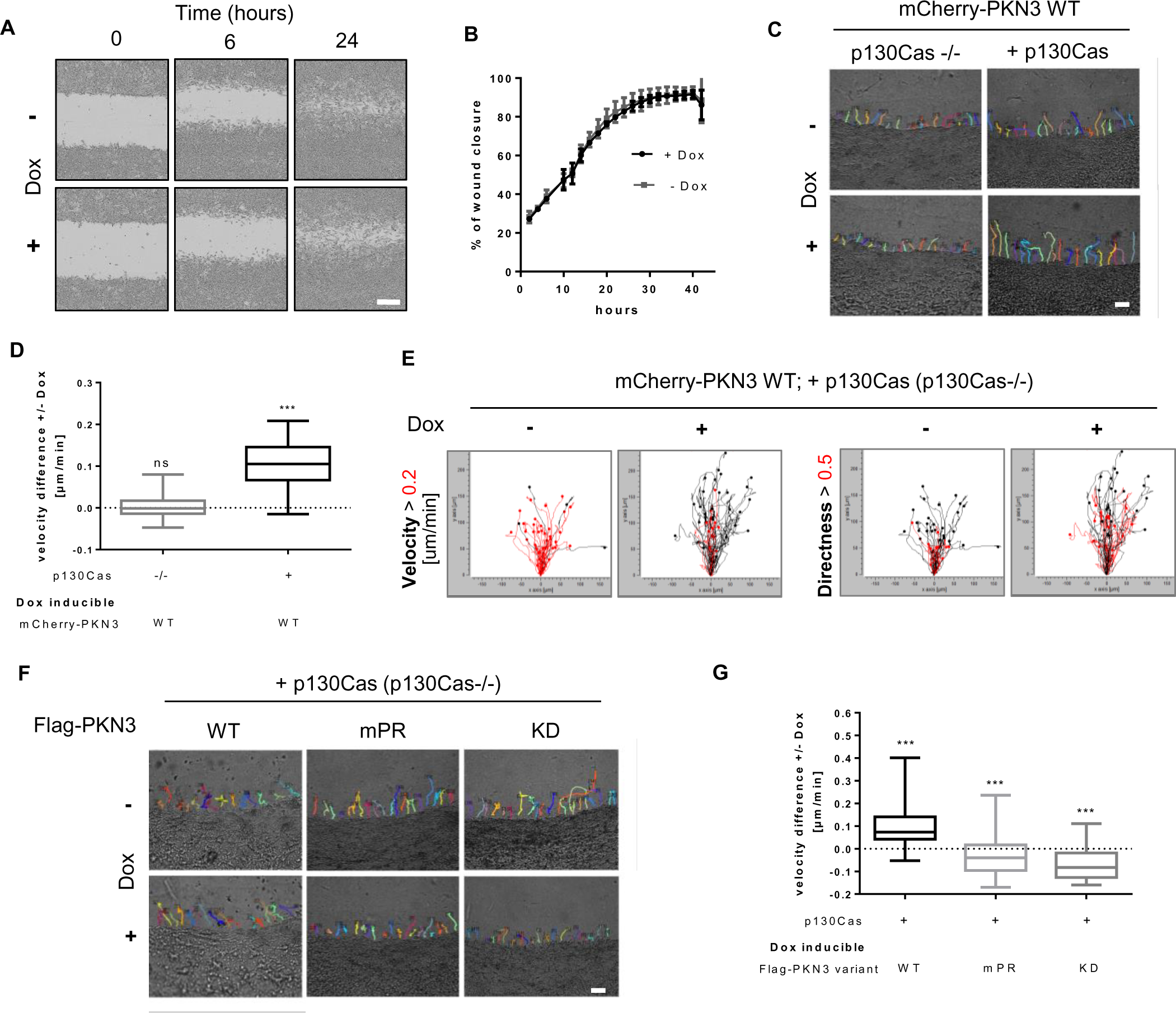
The interaction of p130Cas with PKN3 is required for PKN3-dependent increase of invasiveness but not 2D migration. **A-B)** 2D migration analyzed by wound healing assay. **A)** Representative images of MEFs p130Cas−/− re-expressing p130Cas with or without 24 h pre-induced for Flag-PKN3 expression at the time of wounding (0 h), after 6 and 24 h are shown (scale bar 300 µm). **B)** Representative graph. **C-G)** Cell invasiveness analyzed by 3D cell-zone exclusion assay. Cell migration into collagen was recorded using time-lapse video microscopy with frames being collected every 5 min for 18 h, starting 2 h after collagen scratch, with or without Dox supplementation. Expression of different constructs was pre-induced day before. **C)** Representative tracking maps of cells (MEFs p130Cas−/− or MEFs p130Cas−/− re-expressing p130Cas) migrating into collagen supplemented with or without Dox for induction of mCherry-PKN3 expression (scale bar 100 µm). Relevant supplemental movies S4-5 are included. **D)** Quantification of cell migration velocity difference induced by Dox-inducible mCherry-PKN3 expression in the p130Cas null background with or without re-expression of p130Cas. **E)** Migration maps comparing the velocity and directionality of cells described in D). The boxed traces represent the length (above 0.2 µm black, below red) and directionality (above 0.5 black, below red) of cell movement of each individually tracked cell plotted from the same point of origin. **F)** Representative tracking maps of MEFs p130Cas−/− re-expressing p130Cas migrating into collagen supplemented with or without Dox for induction of Flag-PKN3 variant expression as indicated (scale bar 100 µm) and their quantification **G).** Box and whisker graphs show quantification of cell migration into collagen out of 3 independent experiments (n=60 cells). Statistical significance comparing induced and non-induced cells was evaluated by one-way ANOVA or oneway ANOVA on ranks followed by Turkey post-hoc test (***P < 0.001).

Next, we investigated the potential role of PKN3 for invasive cell migration in 3D collagen. To analyze the effect of PKN3 on cell invasiveness, we utilized a 3D cell-zone exclusion assay as characterized in Van Troys et al. (Van Troys et al., 2018). In this assay, a monolayer of cells grown on a thick layer of collagen is wounded and then the system is overlaid with another layer of collagen. The cells migrating into the wound adopt the morphology and mechanisms of movement of cells migrating in 3D but can be monitored in one focal plane and their movement can be easily tracked and analyzed as in 2D (details described in Methods). p130Cas−/− MEFs re-expressing p130Cas clearly moved faster and more individually compared to p130Cas−/− MEFs (Fig. 6, C and D, and Movie S4 versus 5). Induced expression of PKN3 in p130Cas−/− MEFs (Fig. 6, C and D, and Movie S5) was not sufficient to increase cell migration or to promote individual cell movement. In contrast, activation of PKN3 expression in p130Cas−/− MEFs re-expressing p130Cas led to an increase of cell speed and invading distance compared to those of non-induced cells but did not change cell directionality (Fig. 6, C-E, and Movie S4). Notably, induced expression of PKN3 mPR and KD variants did not induce invasiveness of p130Cas−/− MEFs re-expressing p130Cas (Fig. 6, F-G) but rather had the opposite effect, suggesting that PKN3-p130Cas interaction and PKN3 activity are important for PKN3-induced cell invasiveness of MEFs in collagen.

To test whether PKN3 promotes the invasiveness of MEFs by phosphorylation of the p130Cas SRD domain, we performed the 3D cell-zone exclusion assay with p130Cas−/− MEFs re-expressing GFP-p130Cas variants (WT, 15AN) or GFP as control with or without Dox to induce mCherry-PKN3. Re-expression of GFP-p130Cas (WT or 15AN) caused an increase of cell invasive capability in 3D collagen (Fig. S4 C). Cell treatment by Dox further slightly increased the migration velocity of cells re-expressing both GFP-p130Cas WT and GFP-p130Cas 15AN in collagen compared to that of non-induced controls (Fig. S4, C and D), indicating that phosphorylation of p130Cas in the SRD domain by PKN3 is not involved in the PKN3-dependent increase of MEF invasiveness. Taken together, our results suggest that PKN3 expression increases cell invasiveness of MEFs and that this induced invasiveness is dependent on p130Cas. Furthermore, the PKN3 activity and its ability to interact with p130Cas is required for PKN3-induced cell invasiveness of MEFs, but is independent of PKN3-mediated phosphorylation of the p130Cas SRD domain.

### PKN3 regulates growth and invasiveness of Src-transformed MEFs in a 3D environment through interaction with p130Cas

Having established the importance of the PKN3–p130Cas interaction for cell proliferation and cell migration in a 3D environment, we next analyzed whether this interaction also influences the growth and invasive behavior of SrcF-transformed cells. We first analyzed the effect of SrcF-induced transformation on the level of endogenous PKN3. We found that PKN3 expression is significantly increased in MEFs transformed by activated Src (almost 2×; *p* < 0.009) and that this increase is dependent on the presence of p130Cas (Figs. 7 A and S5 A). To reduce the contribution of endogenous PKN3 in subsequent experiments, we inactivated PKN3 in SrcF p130Cas−/− MEFs re-expressing p130Cas (SC cells) using CRISPR/CAS9 system (Methods; Fig. 7 B). All PKN3 KO clones (SCpkn3−/−) tested exhibited significantly reduced proliferation compared to that of the parental cells (Fig. 7 C). Consistent with this, reintroduction of PKN3 (mCherry-Flag fused), but not PKN3 mPR or mCherry alone, under a Dox-controlled promoter (Fig. 7 D) led to rescue of cell growth in SCpkn3−/− cells when Dox was supplemented to the medium (Figs. 7 E and S5 B).

**Figure 7.**
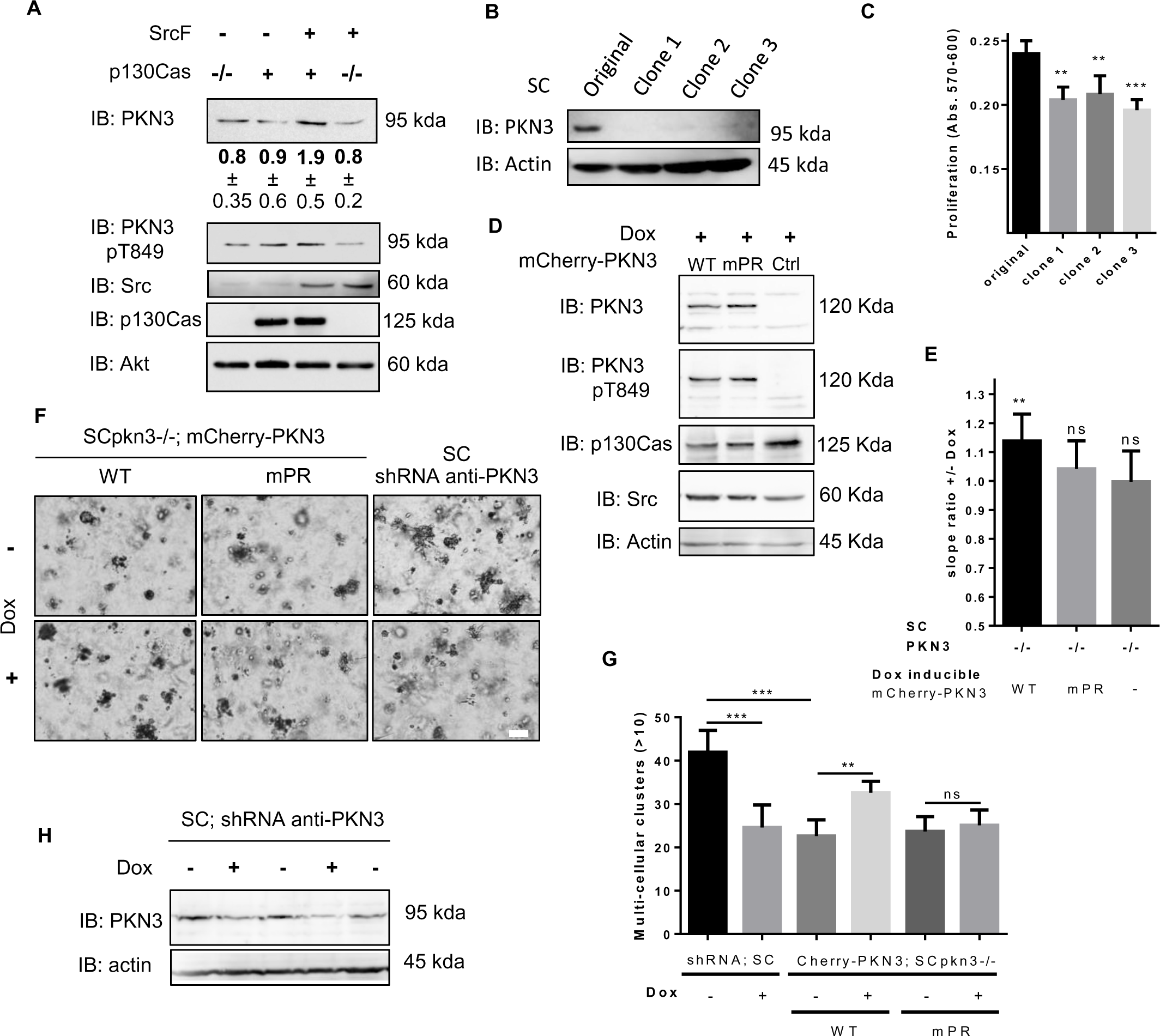
PKN3 regulates growth in 2D and in 3D environment of Src-transformed MEFs through interaction with p130Cas. **A)** Immunoblotted lysates from MEFs p130Cas−/− re-expressing p130Cas with or without transformation by constitutively active Src (SrcF, SC cells). Src and p130Cas were detected by corresponding antibodies anti-Src or anti-p130Cas, respectively. Antibodies anti-PKN3 or anti-pT849 PKN3 were used for PKN3 and anti-Akt served as loading control. Densitometric quantification of protein PKN3 amount is indicated below. **B)** Immunoblot of SC cells and SC cells with *PKN3* gene inactivated using CRISPR/CAS9 (SCpkn3−/−). Inactivation of PKN3 expression is visualized by antibody anti-PKN3. Antibody anti-actin was used as loading control. **C)** Quantification of SC cells proliferation rate with or without inactivated *PKN3* gene cassette by AlamarBlue method (72 h after cell seeding). The graph shows mean absorbance measurement at 570 nm with reference at 600 nm corrected for the initial deviations of cell seeding counts. **D)** Immunoblotted lysates from SCpkn3−/− cells treated by Dox to induce expression of mCherry-PKN3, mCherry-PKN3 mPR or mCherry alone. PKN3 constructs were detected by anti-PKN3 and anti-pT849 PKN3antibodies. Src and p130Cas were detected by anti-Src or anti-p13Cas antibodies, respectively. Antibody anti-actin served as loading control. **E)** Quantification of cell growth change induced by Dox-inducible mCherry, mCherry-PKN3 WT or mCherry-PKN3 mPR expression in SCpkn3−/− cells measured by xCELLigence RTCA system. Slope ratios reflecting cell growth were calculated from the log growth phase of cell growth (representative curves are shown in supplementary figure S5B). **F)** SC and SCpkn3−/− cells stably expressing mCherry-PKN3 WT, mPR or shRNA anti-PKN3 in Dox-dependent manner were embedded into Matrigel. Photographs were taken on day 7, scale bar represents 100 µm; enlarged image for each sample is shown with subtracted background using ImageJ. **G)** Multi-cellular clusters (>10 cells) indicating proliferating cells were quantified relative to individual cells or small cell aggregates on day 7. Average mean is shown. **H)** Immunoblot analysis showing effectiveness of PKN3 knockdown in SC cells. Antibody anti-actin was used as loading control. All error bars indicate means ± SD calculated from 3-5 independent experiments (each in triplicates C), E); in duplicates G)). Statistical significance comparing induced and non-induced cells (E, G) was evaluated by one-way repeated ANOVA followed by Turkey post-hoc test (**P < 0.01), among variants/groups (C, G) by one-way ANOVA followed by Turkey post-hoc test (**P < 0.01; ***P < 0.001).

To mimic the tumor environment, we further examined the growth of SCpkn3−/− cell lines in a 3D environment, similarly as reported in Unsal-Kacmaz et al. (Unsal-Kacmaz et al., 2012). Induced expression of PKN3 in SCpkn3−/− cells promoted a more aggressive behavior characterized by the appearance of a larger number of multicellular clusters in 3D Matrigel when compared to that of uninduced cells (Fig. 7, F and G). In contrast, inducible expression of PKN3 mPR did not significantly change cell growth. To further confirm our results that PKN3 supports aggressive cell behavior of SrcF-transformed MEFs in Matrigel, we also established stable SC cell line inducibly expressing shRNA anti-endogenous PKN3 (Fig. 7, F-H). Consistent with the previous findings, the enhanced growth of parental SC cells compared to SCpkn3−/− cells was reduced by Dox-induced shRNA anti-PKN3 to a basal level of SCpkn3−/−cells (Fig. 7 G). Taken together, these results suggested that PKN3 supports a more malignant phenotype in SrcF MEFs and that interaction of PKN3 with p130Cas may at least partly mediate this effect.

p130Cas has been previously shown to be crucial for invasion of SrcF-transformed MEFs (Brábek et al., 2004). To investigate whether PKN3 could influence the migration of SrcF-transformed MEFs in a 3D environment, we used the 3D cell-zone exclusion assay. In agreement with the results obtained in untransformed MEFs, in SCpkn3−/− cells, induction of PKN3 but not PKN3 mPR expression led to an increase of cell invasion (Fig. 8, A and B). Consistent with this, inducible downregulation of PKN3 in parental SC cells by shRNA led to a decrease of cell migration in collagen (Fig. 8, A and B).

**Figure 8.**
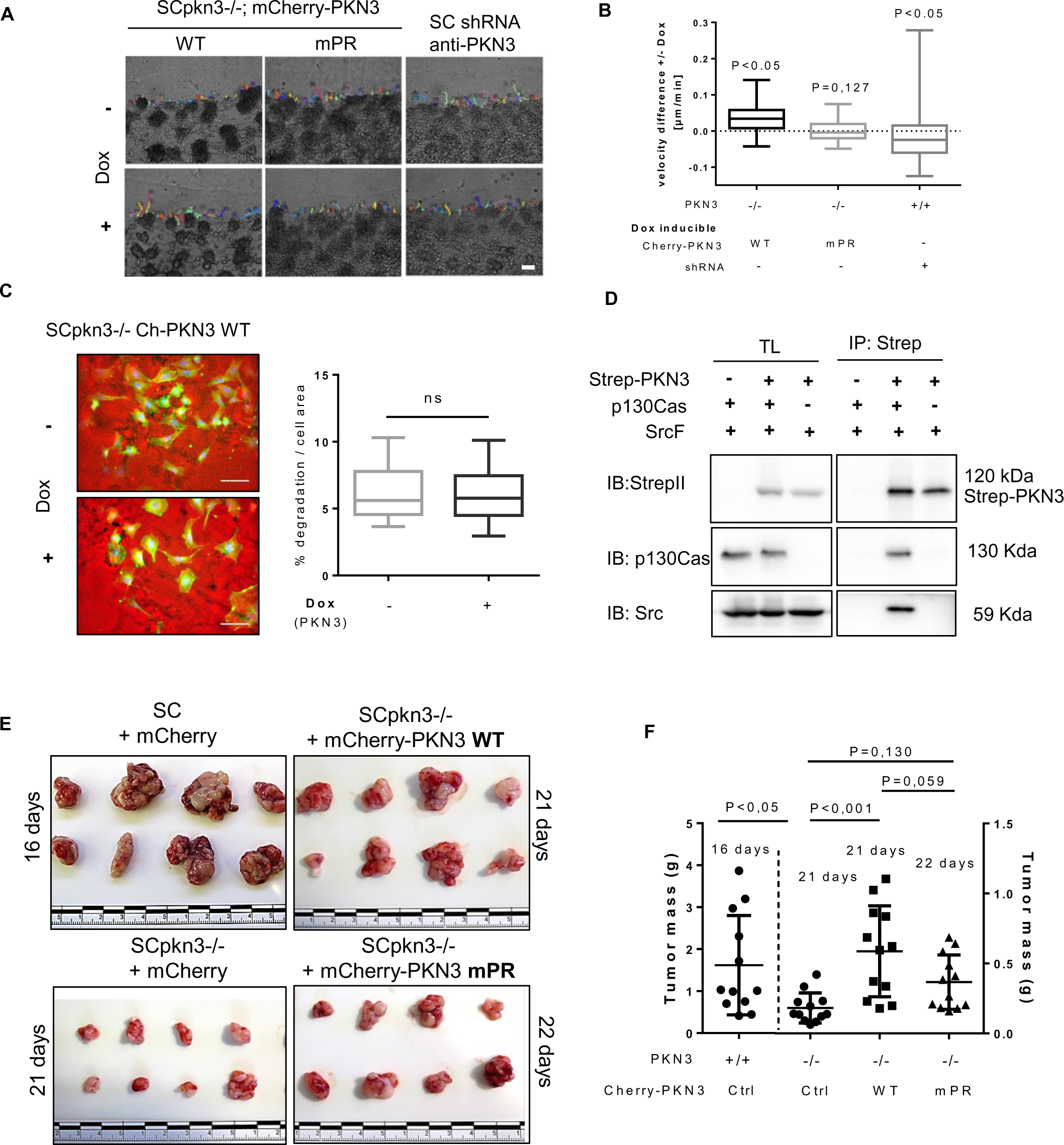
PKN3 regulates cell migration in 3D environment of SrcF-transformed MEFs and tumor growth in vivo through interaction with p130Cas. **A)** Representative tracking maps of SC or SCpkn3−/− cells stably expressing mCherry-PKN3 WT, mPR or shRNA anti-PKN3 in Dox-dependent manner as indicated are shown (scale bar 100 µm). Cells migrated into collagen with or without Dox supplementation. **B)** Quantification of cell velocity difference between induced and non-induced controls. Statistical significance was evaluated from 3-5 independent experiments by one-way ANOVA on ranks followed by Turkey post-hoc test. **C)** Gelatin degradation (Cy3-labeled) of SCpkn3−/− cells expressing Dox-inducible mCherry-PKN3. Representative images are shown (on the left). Scale bar represents 100 µm. Quantification calculated as % degraded area related to total cell area stained by phalloidin 488 (on the right). **D)** Association of PKN3 with Src in p130Cas-dependent manner is shown by co-immunoprecipitations. SrcF-transformed cells with or without p130Cas were transfected by Strep-PKN3 and PKN3 was precipitated by Strep-Tactin^®^ Superflow^®^ resin followed by western blotting and detection by anti-StrepII antibody. Coimmunoprecipitated p130Cas and Src were detected using appropriate antibodies (anti-p130Cas, anti-Src). **E)** 8 representative tumors of SC or SCpkn3−/− cells with Dox-induced mCherry, mCherry-PKN3 or mCherry-PKN3 mPR are shown with the same scale (units in cm). **F)** Their respective quantification expressed as tumor mass weight at defined timescale as indicated. Statistics were evaluated by ANOVA on Ranks Dunn’s Method. TL: total cell lysate; IP: immunoprecipitation; Ctrl: control sample prepared from SC cells.

p130Cas and Src are critical components of podosome-like structures, which are involved in degradation of the ECM (Brábek et al., 2004). As we had shown that PKN3 is also localized to podosome-like structures, we next tested whether the increased invasiveness of PKN3 overexpressing cells was due to their elevated ECM degradation activity. We analyzed the podosome-associated proteolytic activity by gelatin degradation assay and found that gelatin degradation in SCpkn3−/− cells is not affected by the expression of PKN3 (Fig. 8 C). These results indicated that the induced invasiveness of SCpkn3−/− cells expressing PKN3 was reflective of a PKN3-induced migratory phenotype rather than increased degradation of ECM.

Nevertheless, PKN3 was recently demonstrated to be necessary for the bone degradation/resorption activity of osteoclasts, similar to that found for p130Cas (Nagai et al., 2013; Uehara et al., 2017). In both cases, an intact polyproline sequence of PKN3 and the p130Cas SH3 domain were required, as we had validated for PKN3-p130Cas binding, and both sequences were implicated in osteoclast function through an association with Src. We thus hypothesized that the Src-PKN3 interaction is indirect and mediated by p130Cas and tested this by co-immunoprecipitation experiments from SrcF-transformed p130Cas−/− MEFs with or without p130Cas re-expression. As predicted, we found that Src failed to co-precipitate with PKN3 in the absence of p130Cas, confirming that the Src-PKN3 interaction is indirect (Fig. 8 D).

### Interaction of p130Cas with PKN3 is required for PKN3 dependent tumor growth in vivo

As both PKN3 and p130Cas enhance cell proliferation in vitro and can contribute to various stages of tumor development in vivo (Tornillo et al., 2014; Unsal-Kacmaz et al., 2011; Leenders et al., 2004) and to expand our results showing in vitro functional relevance of the PKN3–p130Cas interaction to in vivo analysis, we subcutaneously injected SCpkn3−/− cells stably engineered for inducible expression of either PKN3 WT or PKN3 mPR fused to mCherry, or mCherry alone, into nu/nu mice (total 48). As a positive control, a parental line of SC cells inducibly expressing mCherry was used. The animals were split into four groups (12 per group), all treated with Dox to induce the expression of mCherry-fused PKN3 variants or mCherry alone. The primary tumors were surgically removed at days 21 and 22 after inoculation for the SCpkn3−/− cell lines (+PKN3 WT, +mCherry, and +PKN3 mPR, respectively) and at day 16 in parental SC cells inducibly expressing mCherry owing to large tumor sizes (Fig. 8, E and F). The tumors from parental SC cells with inducible mCherry reached the highest weight despite the shorter period of time. The SCpkn3−/− cells expressing mCherry induced the smallest tumors among all groups. Re-expression of PKN3 WT, but not PKN3 mpR, in SCpkn3−/− cells led to significant increase of tumors weight when compared to unmodified SCpkn3−/− cells (Fig. 8, C and D). Even though the tumors from SCpkn3−/− cells re-expressing PKN3 mPR were removed one day later than those from SCpkn3−/− re-expressing PKN3 WT, their statistical comparison (ANOVA on ranks, Tukey’s post hoc test) yielded an almost significant *p* value of 0.059, indicating that PKN3 likely promotes tumor growth from SCpkn3−/− cells and that the PKN3–p130Cas interaction may be important in this promotion.

We further analyzed individual tumors for expression of mCherry as a reporter for Dox induced expression in the tumor cells using dot blot analysis (Fig. S5 C). Despite the comparable PKN3 mPR and WT protein level produced by Dox-induced cells in vitro, the majority of mCherry-PKN3 mPR expressing tumors exhibited significantly increased mCherry signal compared to that in tumors with mCherry-PKN3 WT. This increase, however, did not lead to comparable tumor growth.

As p130Cas (Brábek et al., 2005) as well as PKN3 (Unsal-Kacmaz et al., 2011) expression is able to promote metastasis, we also evaluated the presence of lung metastases after 14 days post-surgery in animals injected with the parental SC cells, and after 21 days for all SCpkn3−/− cells, as schematically represented in Fig. 5 D. Metastases were clearly observed only in 3 out of 5 surviving mice injected with the parental SC cell line (Fig. S5 E), suggesting that 21 days post-surgery (total 42 days) was not a sufficient time period for the formation of detectable metastases in mice injected with SCpkn3−/− cells.

## Discussion

In this study, we have, for the first time, shown and characterized PKN3–p130Cas interaction and provided new insight into the molecular mechanism of the regulation of tumor-promoting kinase PKN3 and p130Cas-mediated signaling. In particular, we demonstrated that PKN3-p130Cas interaction is necessary for PKN3 to promote malignant growth and invasiveness of SrcF-transformed MEFs. The interaction between mouse p130Cas and mouse PKN3 is mediated by binding of the p130Cas SH3 domain and PKN3 central polyproline region (P_500_PPKPPRL; Fig. 1, A-F). The dynamic of this interaction is potentially regulated similarly as that of other p130Cas SH3 ligands by Src-dependent phosphorylation of the p130Cas SH3 domain on Tyr12, which in its non-phosphorylated state contributes significantly to p130Cas SH3 ligand binding (Gemperle et al., 2017).

The PKN3-p130Cas interaction represents new crosstalk within PI3K–integrin signaling. p130Cas was previously shown to be in complex with PI3K to support anti-estrogenic drug-resistant cell growth (Cabodi et al., 2004). The connection of PKN3 to integrin signaling was indicated by a study showing that PKN3 KO decreased chemotaxis of MEFs toward FN and caused a glycosylation defect of integrin β1 and integrin α5 (Mukai et al., 2016). Moreover, PKN3 promotes osteoclasts to resorb bone matrix via stimulation of integrin effector kinase-Src kinase (Uehara et al., 2017). This effect was strongly dependent on the PKN3 polyproline region that we showed to be responsible for p130Cas binding. Here, we were able to demonstrate that the PKN3-Src association is indeed mediated by p130Cas (Fig. 8 D). Furthermore, consistent with the effect in osteoclasts, we observed a slight activity increase of integrin effector kinase-Src kinase induced by PKN3 overexpression in MEFs (Figs. 7 A, and S4 A and B).

We found PKN3 to co-localize with p130Cas in pro-invasive structures: lamellipodia of MEFs and podosome rosettes of Src-transformed MEFs (Fig. 2, A and B). Notably, PKN3 was not detected at focal adhesions, which are highly redundant in terms of protein composition with podosome rosettes, whereas p130Cas is present in both structures. As PKN family kinases are implicated in cell migration, we investigated the role of PKN3 in cell migration and invasiveness. MEF migration as measured by wound healing assay (2D) was not affected by PKN3 overexpression (Fig. 6, A and B). In contrast, we observed increased cell velocity in collagen (3D) upon induction of PKN3 expression, but not expression of a kinase inactive PKN3 variant (KD) or a PKN3 variant unable to bind p130Cas (mPR). Moreover, the effect of PKN3 expression on migration in 3D was dependent on p130Cas expression (Figs. 6 C-G and 8, A and B). Our results are consistent with independent studies wherein the invasiveness of SrcF-transformed MEFs through Matrigel was greatly decreased in the absence of p130Cas (Brábek et al., 2005) and where MEF PKN3 KO cells exhibited lower migratory activity induced by various treatments than that of MEFs with endogenous PKN3 (Mukai et al., 2016). Similarly, Leenders et al. indicated that PC3 cells with downregulated PKN3 failed to migrate to form network-like structures on Matrigel (Leenders et al., 2004). Although we have not elucidated the exact molecular mechanisms by which PKN3 promotes cell migration, we observed PKN3 binding to actin in a p130Cas-dependent manner (Fig. S5 F) and temporal colocalization of PKN3 with stress fibers (Figs. 3 A and S1, B and C). Given the binding of PKN3 also to regulators of actin dynamics-Rho GTPases (Unsal-Kacmaz et al., 2011), we predict that PKN3 affects intracellular actin dynamics with the co-involvement of p130Cas.

PKN3 is the first kinase documented to phosphorylate p130Cas on Ser/Thr residues (Fig. 4 E), which are abundantly present on p130Cas and yield a dynamic phosphorylation pattern (Makkinje et al., 2009). One of these p130Cas residues, Ser432 (homologous to human Ser428), is abundantly phosphorylated in breast carcinoma tumors (Fig. 4 D) and represents one of the major Ser/Thr phosphorylation sites in MEFs cultivated in vitro (Fig. 4 F). Notably, some phosphoproteomic studies have shown that this phosphorylation is enriched in response to insulin, which is also a potent activator for PKN3 (PhosphoSite database, Leenders et al., 2004)), and correlates with PKN3 activity in breast carcinoma tumors (Fig. 4 D). The effects of this phosphorylation as well as that of p130Cas in the SRD domain are still unknown, as p130Cas−/− MEFs with reintroduced GFP-p130Cas 15AN increased cell growth and invasiveness similarly as GFP-p130Cas WT in a PKN3-dependent manner (Figs. S3 D, S4 C and D).

Notably, a p130Cas paralog termed Nedd9 was found to be phosphorylated on Ser369, which is homologous to p130Cas Ser432. Mutation of Ser369 to alanine, similarly to mutation of p130Cas Ser432, induces p130Cas mobility shift upon SDS-PAGE from 4 bands to two faster migrating bands. Although the kinase responsible for Nedd9 Ser369 phosphorylation is unknown, this modification was shown to induce Nedd9 proteasomal degradation (Hivert et al., 2009). Protein p130Cas, however, is stable during the cell cycle and exhibits a constant expression pattern in cells. Therefore, the effect of p130Cas Ser432 phosphorylation is likely different from that of Ser369 in Nedd9.

Furthermore, transformation of MEFs by a constitutive active Src led to an increase of PKN3 endogenous protein levels in cells grown on plastic, which was dependent on the presence of p130Cas (Figs. 7 A and S5 A). Similarly, PKN3 levels were shown to be elevated by its activators PI3K (Leenders et al., 2004) and RhoA/C (Unsal-Kacmaz et al., 2011), and by the oncogenic form of Ras (Leenders et al., 2004). This suggests that PKN3 functions as an effector of various signal transduction pathways that mediate cell growth and transformation.

Several studies have presented evidence for the significant contribution of PKN3 and p130Cas to tumor growth as well as metastasis formation (Kang et al., 2015; Brábek et al., 2005; Unsal-Kacmaz et al., 2011; Leenders et al., 2004; Tornillo et al., 2014). Our data support these results and furthermore suggest that binding of PKN3 to p130Cas is important for PKN3-induced tumor growth. We hypothesize that targeting p130Cas, PKN3, and their co-operation may be exploited in cancer therapy and, based on our insight in PKN3 signaling in osteoclasts, potentially in pathophysiological signaling related to bone disease such as osteoporosis, rheumatoid arthritis, and periodontal disease as well. Moreover, a therapeutic agent that targets PKN3 by RNA interference (siRNA Atu027) has already finished Phase I clinical trials in a study with patients exhibiting different solid advanced and metastatic tumors, showing very promising results without any adverse effects (Schultheis et al., 2014), further supporting the validity of this strategy.

## Material and methods

### Antibodies and reagents

The following antibodies were used: Akt (rabbit pAb, #9272S), phospho-Akt (Ser473, rabbit mAb D9E, #4060S), phospho-p130Cas (Tyr165, rabbit pAb), p44/42 MAPK (i.e., Erk1/2, rabbit mAb 137F5, #4695), phospho-Src family (Tyr416, rabbit pAb, #2101S), Stat3 (rabbit mAb D3Z2G, #12640), Phospho-Stat3 (Tyr705, mouse mAb 3E2, #9138S), Phospho-Myosin Light Chain 2 (Ser19, mouse mAb, #3675), Phospho-S6 Ribosomal Protein (Ser235/236, rabbit mAb 2F9, #4856), Phospho-GSK-3α/β (Ser21/9, Rabbit mAb 37F11, #9327) (Cell Signaling Technology); p130Cas (mouse mAb 24), Paxillin (mAb 349) (BD Transduction Laboratories); Src (Calbiochem, mouse mAb 327, Ab-1); FAK (rabbit pAb, C–20), Actin (goat pAb, C–11) (Santa Cruz Biotechnology); MAPK (i.e., Erk1/2, mouse mAb V114A, Promega); GST (rabbit pAb, G7781; Sigma Aldrich); PKN3 (rabbit pAb, NBP-130102), mCherry (rabbit pAb, NBP2-25157), StrepII (mAb 517, NBP2-43735) (NOVUS Biologicals); PKN3 (rabbit pAb, AP14628A; Abgent); and phospho-PKN3 (human Thr860 = mouse Thr849) (Pfizer, Oncology). Immunoprecipitations were carried out using a Flag antibody (mouse mAb M2, Sigma Aldrich) and GFP antibody (Abcam, rabbit polyclonal ab290). Secondary antibodies fused to HRP (Abcam) were used as recommended by the manufacturer. The secondary antibodies for fluorescence imaging were: anti-rabbit (Alexa-546) and anti-mouse (Alexa-594, Alexa-633; Molecular Probes).

The reagents used were doxycycline hydrochloride, blasticidin, and puromycin (Sigma Aldrich), glutathione (reduced, LOBA Chemie), Phalloidin 405 (Thermo Fisher Scientific), FN (Invitrogen), collagen R solution (SERVA), Matrigel (Corning), Strep-Tactin^®^ Superflow^®^ resin (IBA Lifesciences), nProtein A Sepharose 4 Fast Flow (GE Healthcare), anti-Flag M2 affinity resin (Sigma), and p-nitrobensyl mesylate (Abcam).

### Cell lines and culture conditions (shRNAs, transfection, retrovirus, and lentivirus preparation)

p130Cas−/− MEFs were obtained from Steven Hanks (Vanderbilt University, Nashville). p130Cas−/− MEFs expressing constitutively active mouse Src Y527F (SrcF cells) or both p130Cas WT and Y527F (SC cells) were prepared using the LZRS-MS-IRES-GFP retroviral vector and the Phoenix E packaging line as described previously (Brábek et al., 2004).

To generate SC PKN3 null, cells were transiently transfected by sgRNA anti-PKN3 (sequence provided in Table S2) and a puromycin resistance-expressing CRISP-CAS9 plasmid. Cells were selected by puromycin for 2–3 days followed by clonal isolation. Success of CRISPR targeting was analyzed by PCR and Surveyor nuclease assay as published (Ran et al., 2013) and by western blots. Sequences around the CRISPR modification site were amplified by PCR from three selected mammalian clones, cloned into pJET1.2/blunt using a CloneJET PCR Cloning Kit (ThermoFisher), and sequenced (at least six from each). Sequencing revealed that CRISPR nicking most frequently caused frameshift -1 bp or + 49/+37 bp. Potential off-targets were predicted (http://tools.genome-engineering.org) and the three most potent tested by Surveyor assay (results negative, primer sequences provided in Table S2). SC cells with Dox-inducible PKN3 shRNA (see plasmid construction, below) were prepared similarly as described previously (Leenders et al., 2004; Czauderna et al., 2003).

To stably deliver Flag-fused or mCherry-Flag-fused constructs, retroviral pMSCV-puro or lentiviral pLVX-Tet-On Advanced system (Clontech) was used. In brief, cells were infected with viral supernatant generated in transfected Phoenix E packaging cells (pMSCV system) or produced in HEK293T cells co-transfected by pVSV-G and psPAX2 plasmids together with pLVX-Tet-On in the first round or with pLVX-tight-puro in the second. All cells were then selected with puromycin (2.5 µg/ml) or/and blasticidin (2–4 µg/ml) and alternatively sorted by FACS (1–3 rounds) after transient (24 h) induced expression by Dox. Cells without fluorescent protein were cloned and tested by western blot. SCpkn3−/− cells from clone 1 were used for reintroduction of PKN3 (mCherry-Flag fused constructs).

To re-introduce GFP fused p130Cas variants to p130Cas−/− MEFs, a retroviral pBabe system was used. After infection of p130Cas−/− MEFs by viral supernatants prepared in HEK293T cells co-transfected by pVSV-G, Gag-Pol, and pBabe plasmids, respectively, GFP positive cells were selected by FACS.

MDA-MB-231 cells were provided by Dr. Marie Zadinova from Charles University. All transfections were carried out according to the manufacturer’s protocol using Jet Prime (Polyplus Transfection) or PEI transfection reagent (Polysciences). All cells were cultivated in full DMEM (Sigma) with 4.5 g/l L-glucose, L-glutamine, and pyruvate, supplemented with 10% FBS (Sigma) and ciprofloxacin (0.01 mg/ml) at 37 °C and 5% CO2. Cells were tested for mycoplasma contamination by PCR.

### Plasmid construction

cDNAs coding for p130Cas SRD domain variants (WT, 15AN; sequences provided in Table S1) were commercially synthesized and cloned into the pMA-T vector (geneArt, Life Technologies). The 15AN variant was generated by mutation of all 15 Ser/Thr to Ala or Asn. Mutant variant S432A (KRLS to KRLA) was created subsequently by whole plasmid synthesis using Q5 polymerase (New England Biolabs) and respective site-directed mutagenesis primers (listed in Table S2). Following PCR, 5U of DpnI was added to each reaction and incubated for 1.5 h at 37 °C. Individual mutated clones were screened by sequencing. Sequences of all SRD variants were then switched with the original p130Cas SRD domain cassette within the p130Cas sequence (pUC19 vector) using Bpu10I/SacII sites. pEGFP-C1 p130Cas variants and pGEX-p130Cas-SH3 domain constructs were prepared similarly as described previously (Janoštiak et al., 2011; Braniš et al., 2017). GFP-fused p130Cas variants were introduced into the pBABE retroviral expression vector by EcoRI and blunt end (generated by BamHI/AgeI restriction followed by fill in by Klenow) cloning. All constructs were verified by sequencing. To prepare the GSK3-derived peptide fused to GST, a pair of phosphorylated oligonucleotides (see Table S2) was annealed and inserted in frame at the 3’ end of the GST (pGEX vector) using BamHI/EcoRI sites. Production of GST fused proteins (SH3, GSK3) from the pGEX system were carried out as in Gemperle et al. (Gemperle et al., 2017). Oligos for sgRNA construction and primers for CRISPR off-target screening were designed and plasmids constructed (LentiCRISPR, BsmBI site) as published (Ran et al., 2013) and are listed in Table S2. PKN3 shRNA was cloned into a lentivirus-adapted vector system allowing Dox-inducible expression of shRNAs (Czauderna et al., 2003) via PCR followed by ligation and DpnI mediated cleavage of template DNA. Primers are listed in Table S2. Correct insertion of the shRNA sequence was confirmed first by restriction analysis to confirm the reintroduced BsrgI restriction site, and then with DNA sequence analysis.

cDNAs coding for whole mouse PKN3 with added sequence for Flag epitope and shorter sequence with mutations to create mPR and KD variants (sequences provided in Table S1) were commercially synthesized and cloned into pMK-RQ or in pMA-T vectors, respectively (geneArt, Life Technologies). The mPR mutant was created to abrogate binding to p130Cas (motif P_500_PPKPPR to PAPSAPR) similarly as suggested in Gemperle et al. (Gemperle et al., 2017). The KD mutation was designed as published for human PKN3 (Leenders et al., 2004). Sequence for mPR was swapped within whole PKN3 in the pMK-RQ vector using NdeI/HindIII sites, and that for KD by NdeI/BamHI followed by cloning PKN3 variants (WT, mPR, KD) to other corresponding vectors: mCherryC1/eGFP C1 (BsrgI/EcoRI), pMSCV-Puro (XhoI/EcoRI), and pLVX-Tight-Puro (BsrgI/EcoRI). A lentiviral expression vector pLVX-Tight-Puro allowing Dox-inducible gene expression was also prepared with mCherry-fused PKN3 variants via NheI/EcoRI and XbaI/EcoRI sites. Flag-fused mCherry control was prepared similarly, except prior to cloning to the pLVX-Tight-Puro system, Flag sequence was inserted in frame in front of mCherry in mCherry C1 vector by annealing the phosphorylated oligonucleotides described in Table S2 and digested NheI/HindIII vector sites. Prior to kinase reactions in vitro, Flag-PKN3 was also cloned to the StrepII pcDNA3 vector using blunt end ligation (AfeI/EcoRV vector sites, Flag-PKN3 cleaved from pMSCV-puro by BsrgI/EcoRI and filled in by Klenow). CFP or mCherry-fused LifeAct was prepared as published (Riedl et al., 2008). All constructs were verified first by restriction analysis and then by sequencing.

### Preparation of cell extracts and immunoblotting

Cell were lysed in RIPA (total lysates; composition: 150 mM NaCl; 50 mM Tris-HCl, pH 7.4, 1 % Nonidet P-40, 0.1 % SDS, 1 % sodium deoxycholate, 5 mM EDTA, 50 mM NaF) or in 1% Triton lysis buffer (immunoprecipitations, pull downs; composition: 50 mM Tris HCl [pH 7.1], with 150 mM NaCl and 1% TRITON X−100) supplemented with protease (MixM; SERVA) and phosphatase inhibitors (MixII; SERVA) followed by immunoblotting as described previously (Janoštiak et al., 2014). In brief: protein extracts were separated using SDS-PAGE under denaturing conditions (6−15 % gels) and were transferred to nitrocellulose membrane (Bio-Rad Laboratories). Membranes were blocked with 4 % BSA or 3% milk-TBST (Tris-buffered saline and 0.05% Tween 20), incubated with the indicated primary antibodies overnight at 4 °C, and then incubated with HRP-linked secondary antibodies at RT for 1 h, washed extensively in TBST, and developed using an AI600 System (GE Healthcare). To improve the separation of p130Cas by SDS-PAGE, a ratio of acrylamide/bisacrylamide 30:0.2 was used.

Immunoprecipitations (see kinase assays, below), pull-downs, and far western experiments were carried out similarly as in Gemperle et al. and Janoštiak et al. (Gemperle et al., 2017 and Janoštiak et al., 2014). For far-western-blot analysis, the protein blots were incubated with 2 µg/ml recombinant human GST-p130Cas/BCAR1 (Abcam) or purified mouse GST-p130Cas SH3 diluted in 1 % BSA in TBST overnight followed by washing with TBST and incubation (2 h, 4 °C) with anti-GST antibody (Sigma). After extensive washing with TBST, blots were treated with HRP-conjugated secondary antibodies and developed using the AI600 System. Band intensity was determined using Fiji (ImageJ; National Institutes of Health; Schindelin et al., 2012).

### Kinase assays

Cells were transfected with either Flag- or StrepII-fused mouse PKN3 (WT or KD) using the PEI transfection reagent (Polysciences) according to manufacturer instructions. After 48 h, cells were washed with PBS and lysed in standard lysis buffer (50 mM Tris-HCl [pH 7.1], 150 mM NaCl, 1 % Triton-X-100) with protease and phosphatase inhibitors (SERVA) and 10 mM glycerol-2-phosphate (Sigma). Proteins were immunoprecipitated with anti-Flag M2 affinity resin (Sigma) and eluted with Flag peptide as described previously (Unsal-Kacmaz et al., 2011). StrepII-fused PKN3 was precipitated with Strep-Tactin^®^ Superflow^®^ resin (IBA Lifesciences) and eluted with 1× Buffer E (Strep-Tactin Elution Buffer, IBA Lifesciences). Kinase assays were carried out in conditions described previously (Unsal-Kacmaz et al., 2011; Leenders et al., 2004). GFP-fused p130Cas variants (WT, 15AN, S432A; see plasmid construction section, above) were immunoprecipitated from transiently transfected cells using anti-GFP 3E6 antibody (Invitrogen) and nProtein A Sepharose 4 Fast Flow (GE Healthcare). Subsequently, immobilized p130Cas was dephosphorylated using Lambda protein phosphatase (New England Biolabs) as specified by the manufacturer. Phosphatase was inactivated by two washes with TBS containing 10 mM sodium orthovanadate and 20 mM EDTA and two washes with TBS and p130Cas variants were eluted with 0.1 M glycine pH 3.5 for 10 min in RT. After elution, pH was equilibrated by adding the corresponding volume of 1 M Tris pH 9.2 and proteins were used as a substrate in reaction with 1 mM ATPγS (Sigma). After 45 min in 35 °C, reactions were stopped with EDTA to a final concentration of 20 mM and alkylated with 50 mM p-nitrobensyl mesylate (Abcam) at RT for 2 h. Samples were resolved using SDS-PAGE and immunoblotted using anti-thiophosphate ester antibody (clone 51-8, Abcam). GSK3-derived peptide fused to GST was prepared and used as a positive control for PKN3 activity as described previously (Unsal-Kacmaz et al., 2011) and its phosphorylation detected by anti-Phospho-GSK-3α/β antibody. For autoradiography, kinase reactions were performed similarly, with 5 µCi [γ-32P] ATP added to the reaction. The samples were heated for 10 min at 95 °C, resolved using SDS-PAGE, and subjected to autoradiography.

### 2D and 3D migration assays

Scratch-based migration assays in 2D were carried out using an IncuCyte automated imaging system (Essen BioScience). Briefly, MEFs with Dox-inducible PKN3 WT were seeded onto 96-well plates (Corning) at a density of 100,000 cells/well (10% serum in DMEM) and half of the wells were supplemented with Dox (final 250 ng/ml). After 24 h, monolayers of cells were scratched using a scratching apparatus that produced strongly identical scratches in each well. The IncuCyte system was programmed to obtain real-time phase-contrast images of the wounds every 2 h for 2 days. Cell migration was automatically quantified and expressed as relative wound density, which indicates the ratio of sharpness of the wounded area and of the adjacent non-wounded area. The IncuCyte imaging system was then used to automatically calculate the area of each wound at each time point up to the point of complete closure of the wound from an average of the quadruplicate.

3D cell zone exclusion assays were carried out using the JuLI™ Br (NanoEnTek) system with two microscopes situated in the incubator. Collagen R solution (4 mg/ml, SERVA) was diluted to the final mix: 1 mg/ml collagen, 1 % serum (to decrease the effect of proliferation), 1x DMEM, 15 mM HEPES (750 mM), 8.5 mM NaOH (1 M), 0.4 % NaHCO3 (7.5 %, Sigma), and 5 µg/ml folic acid. Next, 40 µl of the collagen mix was then added into each well (total 6) of two 96-well plates and let to polymerize at 37 °C. Cell suspension (100 μl; 1 × 106 cells/ml) non-induced or 24 h Dox-preinduced cells were added on top of the collagen gel. After cell attachment (4 h), the medium was removed and scratch was performed. Immediately after removal of the rest of media in the generated wound, another layer of collagen mix (100 µl) was added (schema of the experiment is shown in Van Troys et al. (Van Troys et al., 2018)). Collagen was allowed to polymerize for 15 min at RT to prevent formation of bubbles in the collagen interface, then for another 15 min at 37 °C. Finally, 100 µl of DMEM with 1 % serum supplemented with or without Dox (500 ng/ml) was added on the top of each well and CellTracker software (Piccinini et al., 2015) was used to manually track at least 60 random (fastest) (top) migrating cells across the cell/collagen interface in the main focal plane from three independent replicates. To create cell migration/tracking maps the “Chemotaxis And Migration Tool” version 2.0 (Ibidi) was used.

### 2D proliferation and 3D cultures in Matrigel

To determine the difference in cell proliferation capacity among individual clones, an AlamarBlue assay was used. For this, 10,000 cells per well were seeded in a 96-well plate and after cell attachment (4 h), medium was removed and replaced by a culture medium solution containing 10 % AlamarBlue. The plates were further incubated for 2 h at 37 °C prior to initial cell mass measurement followed by replacing with standard medium (± Dox if required). Cell growth was than measured again after 72 h using a culture medium solution containing 10 % AlamarBlue to assess the increment of cell mass. Absorbance was measured at 570 nm (with reference at 600 nm) using an Infinite M200 PRO microplate reader. As a control, AlamarBlue was added to the cell growth medium without cells. The assay was performed in triplicates and was repeated three times.

xCELLigence RTCA technology, which provides highly standardized experimental conditions, was used to study cell proliferation. The cells were counted (100,000/ml) and then separated to supplement one part with Dox (final 250 ng/ml) and one part without. Subsequently, 10,000 cells with or without Dox were seeded per well of an xCELLigence E-plate in triplicate, left to sit for 30 min at RT, and then transferred to the xCELLigence RTCA device. Data were collected every 30 min for 48–72 h. All data were recorded using RTCA Software version 2.0, which generated curve-slope values reflecting the speed of cell growth. Cell growth was calculated from the log growth phase of the curve starting 4–9 h post Dox-induction followed by processing in MS Excel and comparison to non-induced controls. E-plates were recycled to remove cells and used repeatedly up to three times.

SC cells were embedded into Matrigel (Corning) as previously published (Unsal-Kacmaz et al., 2011). In brief, cells grown in 6-cm dishes with or without Dox for 24 h were detached from culture dishes and suspended in Matrigel solution (per well: 80 µl of Matrigel and 20 µl of medium with 10 % serum and 20, 000 cells) at 4 °C. Then, 100 µl of this Matrigel-embedded cell suspension was overlaid on 40 µl of Matrigel in a 96-well plate. The plate was allowed to solidify at 37 °C in the incubator. After 30 min, DMEM ± Dox (500 ng/ml) was added and replaced every 2–3 days. After 7 days, multicellular clusters were imaged using differential interference contrast (DIC) microscopy (Nikon-Eclipse TE2000-S) and analyzed from duplicated wells. Quantification was based on at least three independent experiments.

### Microscopy and immunostaining

Confocal images/movies were acquired using a Leica TCS SP2 or SP8 confocal microscope system equipped with a Leica 63×/1.45 oil objective (two HyD detectors and two standard PMT detectors, Leica LAS-AF software) followed by processing in Fiji.

Cells were seeded on cover slips coated with human FN 10 µg/ml (Invitrogen), grown for 24 to 48 h, and subsequently fixed in 4 % PFA, permeabilized in 0.5 % TritonX-100, washed extensively with PBS, and blocked in 3 % BSA. The cells were then sequentially incubated with primary antibody (dilution according to manufacturer protocol) for 2 h, secondary antibody for 60 min, and Alexa Fluor 594 phalloidin (Molecular Probes) for 15 min, with extensive washing between each step. In addition, slides were mounted with Mowiol 4–88 (475904; EMD Millipore) containing 2.5 % 1,4-diazobicyclo-[2.2.2]-octane (D27802; Sigma-Aldrich) in the presence or absence of DAPI for nuclear staining and imaged at RT.

Colocalization analyses of PKN3 with p130Cas and actin in lamellipodia were conducted on live cells co-transfected by mCherry-PKN3 variants (mPR, WT, –), GFP-p130Cas and CFP-fused LifeAct. Alternatively, GFP-PKN3 variants and mCherry-LifeAct were used. The cells were placed on glass-bottom dishes (MatTek,) coated with 10 µg/ml FN, transfected, and cultured for 24 h before the experiment. Movies were acquired of cells in DMEM without phenol red supplemented with 10 % Serum at 37 °C and 5 % CO2 using a Leica TCS SP2 or SP8 confocal microscope (63×/1.45 oil objective). Quantification of mCherry-PKN3 WT, mCherry-PKN3 mPR, and mCherry localization to lamellipodia (lifeActin as marker) was calculated by measuring the signal ratio of mCherry fluorescence in lamellipodium versus in cytosol (5 µm from lamellipodium). A total of 160 measurements (50 living cells) gathered in three independent experiments were performed using Fiji.

Gelatin degradation assay was performed according to the manufacturer instructions (QCM™ Gelatin Invadopodia Assay, Merck Millipore). Cells were seeded on gelatin supplemented with DMEM, 10 % serum with or without Dox, and gelatin degradation was assessed after 48 h using a Nikon-Eclipse TE2000-S (10×/0.25 or 20×/0.40 Nikon objective) followed by processing in Fiji.

Lung metastases on the lung parenchyma were visualized using a Carl Zeiss AxioZoom.V16 fluorescence macroscope followed by processing in Fiji.

### Subcutaneous mouse tumor models

Nu/nu mice (total 48, n = 12 per group), 8-week old, were subcutaneously injected with 1 × 106 GFP-positive SC cells (200 µl of suspension in PBS) that either expressed endogenous PKN3 (group 1) or lacked PKN3 expression (group 2, 3, and 4). These cells expressed mCherry vector (group 1 and 2), mCherry-fused Flag-PKN3 WT (group 3), or mCherry-fused Flag-PKN3 mPR (group 4) induced by doxycycline hydrochloride, which was administered via Dox pellets (200 mg/kg) on the day of cell injection. After 16 (group 1, faster tumor growth) or 21–22 days (within groups 2–4), tumors were surgically removed and their weight was determined, followed by disruption by a tissue tearor in RIPA lysis buffer containing protease inhibitors (Serva) and preclearance by centrifugation at 16,000 *g* at 4 °C for 30 min. Tissue lysates were normalized to GFP level (Infinite M200 PRO) and analyzed by immunoblotting (SDS-PAGE separation or dot blot) as described in Janoštiak et al. (Janoštiak et al., 2014). Post-operation, doxycycline hydrochloride was administered via drinking water at 0.2 mg/ml and supplemented with 1 % sucrose. Animals in group 1 were killed 14 days post-surgery, and in groups 2–4 at 21 days post-surgery, and their lungs were collected. Immediately at sacrifice the right lungs were washed in PBS and viewed under a fluorescence microscope to analyze GFP and mCherry-positive metastases grown on the surface, as well as within the lung parenchyma. Right lungs were fixed for 72 h in 4 % PFA for immunohistochemistry analysis. All experiments (surgery, cell transplantation, measurement of tumor weight) were performed blinded and independently by two researchers, and in compliance with the guidelines of the Ministry of Education, Youth and Sports of CR (institutional approval no.70030/2013-MZE-17214).

### In silico analysis

Data of invasive breast carcinoma (1100 tumors in TCGA, provisional) and prostate adenocarcinoma studies (499 tumors, TCGA, provisional) were retrieved from and analyzed using the cBio Cancer Genomics Portal (cbioportal.org) (Gao et al., 2013) or processed using the SigmaPlot software package (Systat Software, Inc., Point Richmond, CA). The available data on protein phosphorylation levels of p130Cas/BCAR1 were renumbered from isoform 6 (Ser474) to isoform 1 (corresponding to Ser432)

### Statistical analysis

Statistical analyses were performed using the SigmaPlot software package (Systat Software, Inc.). Data with a normal distribution were subjected to one-way ANOVA, whereas data failing the normality test were analyzed by one-way ANOVA on ranks followed by Dunn’s post hoc comparison. All compared groups passed an equal variance test. Where not indicated differently, the same cells treated or not treated by Dox were compared. Graphs were created using GraphPad Prism 6. Data are reported as the means ± SD unless otherwise indicated. Correlations statistics were calculated according to the Spearman’s rank and Pearson correlation methods. A p-value of 0.05 was considered to be a borderline for statistical significance. P-values are indicated in the figure legends.

### Online supplemental material

Fig. S1 shows confirmation of PKN3 kinase inactivity, and another western blot showing PKN3 phosphorylation on Thr849 influenced by p130Cas variant expression and that PKN3 colocalizes with stress fibers. Fig. S2 shows PKN3 autophosphorylation, the phospho-pattern of p130Cas from suspension cells, and co-expression of human p130Cas/BCAR1 and PKN3 in prostate and breast tumors. Fig. S3 shows the influence of PKN3 and GFP-p130Cas variants expression on cell morphology and proliferation. Fig. S4 shows that PKN3 induction does not change the activation status of STAT3, ERK, Akt, MLC, and mTOR signaling (only slight increase in Src activity) in cells growing on plastic and that PKN3-dependent invasiveness is independent of phosphorylation in p130Cas SRD. Fig. S5 shows that PKN3 crosstalk with Src is p130Cas dependent. Furthermore, Fig. S5 also provides additional information relevant to tumors and metastasis. Fig. S6 shows that PKN3 binds strongly to the p130Cas SH3 domain and do not bind to the p130Cas SRD domain. Movies S1–S3 show that only PKN3 WT clearly localizes to lamellipodia of p130Cas−/− MEFs re-expressing p130Cas co-transfected by GFP-PKN3 variants and mCherry-LifeActin. Movies S4–S5 show (3D cell zone exclusion assay) that PKN3 increases cell migration velocity in collagen only in the presence of p130Cas. Tables S1 and S2, included as an Excel files, provide a list of synthetized cDNA sequences (geneArt) or oligonucleotides, respectively, used in this study.

## Acknowledgements

This work was funded by Czech Science Foundation grants 15-17419S and 15-07321S, by the Ministry of Education, Youth and Sports of CR within the LQ1604 National Sustainability Program II (Project BIOCEV-FAR), LQ1601 (Project CEITEC 2020), by the project _”_BIOCEV“ (CZ.1.05/1.1.00/02.0109), and by the Charles University Grant Agency (224217). We acknowledge support by the MEYS CR (CZ.02.1.01/0.0/0.0/16_013/0001775 Czech-BioImaging) and support by the Imaging Methods Core Facility at BIOCEV Institution supported by the Czech-BioImaging large RI project (LM2015062 funded by MEYS CR) for their support with obtaining imaging data presented in this paper. We thank Marie Charvátová for technical help and L. Janečková for help with immunohistochemistry. We acknowledge Pfizer as the source of the materials and thank Dr Keziban Unsal-Kacmaz by providing the anti-phosphoThr849 PKN3 antibody (human Thr860). We would like to thank Editage (www.editage.com) for English language editing.

The authors declare no competing financial interests.

## Author contributions

Conceptualization, J. Gemperle, D. Rosel, and J. Brábek; methodology, J. Gemperle, M. Dibus, and L. Koudelková; investigation, J. Gemperle, M. Dibus, D. Rosel, and J. Brábek; resources, J. Brábek; writing: original draft, J. Gemperle; writing: reviewing, J. Gemperle, M. Dibus, D. Rosel, and J. Brábek; visualization, J. Gemperle; supervision, J. Brábek and D. Rosel; funding acquisition, J. Gemperle, M. Dibus, J. Brábek, and D. Rosel.

## Footnotes

Abbreviations used: BCAR1 = breast cancer anti-estrogen resistance protein 1; Dox = doxycycline; FN = fibronectin; KO = knockout; mPR = mutation P_500_PPKPPRL to PAPSAPRL (mouse PKN3 mutant unable to bind p130Cas); MEF = mouse embryonic fibroblast; PKN3 = protein kinase N3; p130Cas = Crk-associated substrate (size 130 kDa); RTCA = real-time cell analysis; SC = p130Cas−/−; MEFs re-expressing p130Cas transformed by SrcF; SCpkn3−/− = SC PKN3 KO; SrcF = constitutively active Src (Src Y527F).

